# The *Candida auris* Adhesin Scf1 Mediates Broad Host-Protein Recognition through Shared Adhesive Regions

**DOI:** 10.64898/2026.09.04.749504

**Authors:** Pratik Vyas, Jiajun Li, Rosario Vanella, Michael A. Nash

## Abstract

*Candida auris* (*C. auris*) is a rapidly emerging, multidrug-resistant fungal pathogen whose persistence is partly driven by adhesion to host tissues and abiotic surfaces. Surface-colonization factor 1 (Scf1) is a *C. auris*-specific adhesin implicated in surface adhesion, biofilm formation, virulence, and persistence in health care settings. However, the human proteins recognized by Scf1 and the response of these interactions to hydrodynamic flow and shear stress were previously unknown. Using yeast surface display, spinning-disk adhesion and FACS-based binding assays, we show that the N-terminal adhesive domain of Scf1 binds the cell-surface glycoprotein mucin-1 (MUC1) and the extracellular matrix proteins vitronectin (VIT) and fibronectin (FN), while exhibiting no detectable binding to collagen I, collagen IV, or fibrinogen. Hydrodynamic shear stress was found to tune ligand preference, with VIT being preferred at equilibrium and MUC1 being the more strongly bound target under flow. Epitope mapping by deep mutational scanning (DMS) of Scf1 against MUC1, VIT and FN identified shared binding patches consisting of predominantly cationic and aromatic residues involved in recognizing all three ligands. Mutations within these regions reduced binding without compromising yeast display expression levels, indicating that these residues are involved in ligand recognition and that mutations at these positions do not compromise global protein folding stability. These findings establish the Scf1 N-terminal domain as a semi-promiscuous ligand-recognition domain for a structurally diverse set of human proteins and suggest that a mechanosensitive adhesion mechanism contributes to *C. auris* colonization.

## Introduction

*Candida auris (C. auris)* is a rapidly emerging fungal pathogen of global health concern. ^1–5^ First identified in Japan in 2009, *C. auris* has now been reported across more than 80 countries and six continents worldwide. ^6,7^ Reported cases of *C. auris* are rising each year, and many cases remain undetected worldwide due to diagnostic challenges. ^8^ *C. auris* spreads rapidly in healthcare settings, causing outbreaks in hospitals and nursing homes worldwide. ^4,9,10^ Some strains exhibit tolerance to all three major classes of antifungals, rendering these infections extremely difficult to treat with high mortality rates (30-60%) for vulnerable patients. ^5,11,12^ At the heart of *C. auris* persistence is its ability to adhere to biotic and abiotic surfaces and form robust biofilms. These biofilms create drug-tolerant communities that resist disinfectants, persist in hospital environments, and facilitate transmission and recurrent outbreaks. ^7,13–16^

Adhesion in *C. auris* is believed to be facilitated by wide-array of cell-surface glycoproteins termed *adhesins.* ^17–20^ However, the human protein ligands recognized by these adhesins are largely unknown. Among these, Surface-colonization factor 1 (Scf1) stands out as a *C. auris*-specific adhesin that promotes surface attachment and is required for virulence in mouse models. ^19^ Scf1 is a 765 amino acid protein comprising an N-terminal region that contains the putative adhesive domain. Structurally, this domain adopts a *Flo-like* fibronectin type III fold (**Fig. 1A**), a motif commonly found in eukaryotic proteins involved in cell adhesion, differentiation, migration and proliferation. ^19,21,22^ The C-terminal region of Scf1 contains a glycosylphosphatidylinositol (GPI) anchor, a post-translational glycolipid modification widely used in eukaryotes to tether proteins to the cell surface. ^19,23^ Given that surface adhesion is the first step in colonization, identifying the human protein ligands recognized by Scf1 is critical for understanding infection biology, biofilm formation and persistence in host environments.

**Figure 1.**
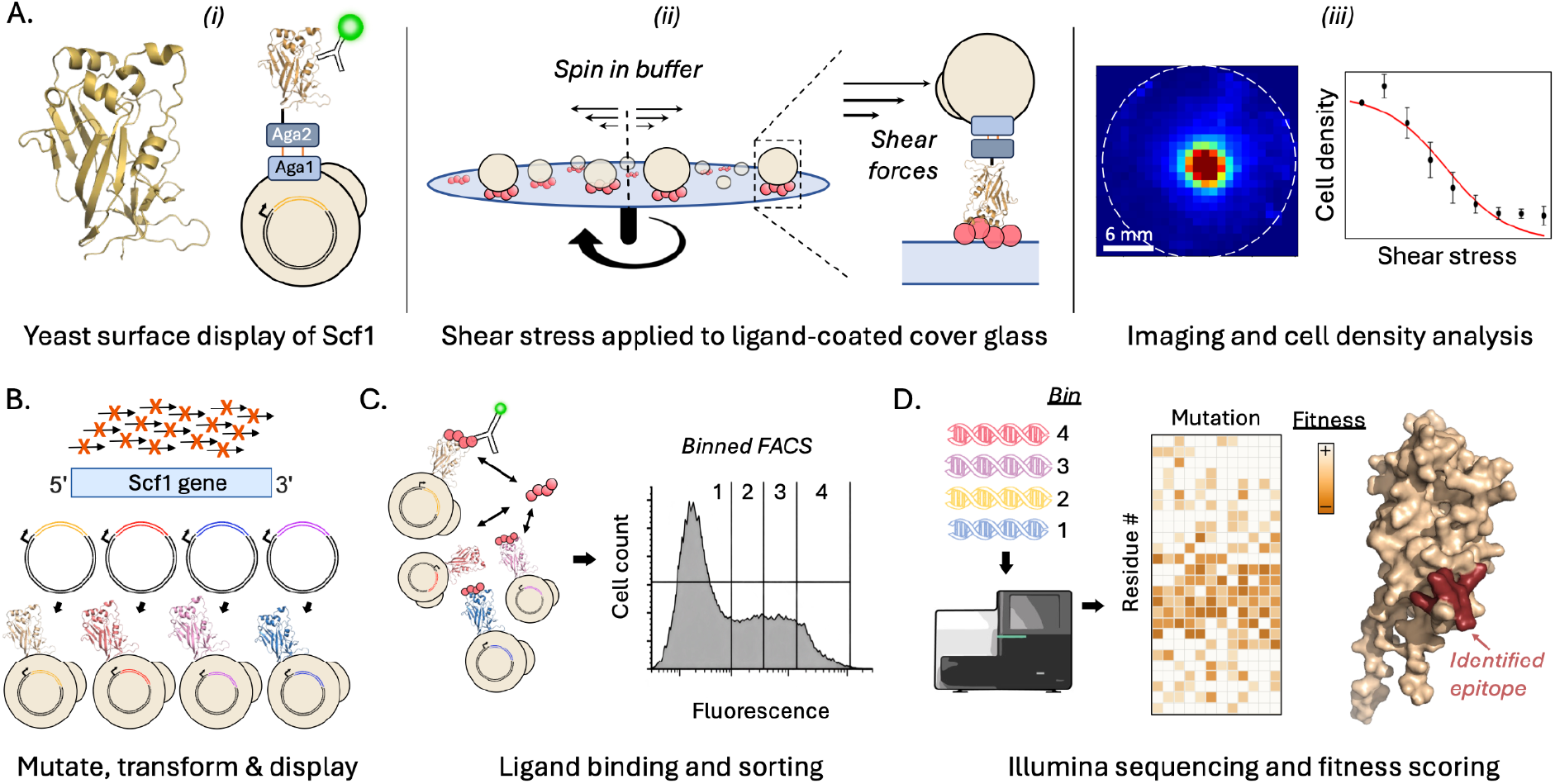
Scf1 ligand-screening and epitope-mapping workflow. **A. Yeast-display and SDA**. (i) AlphaFold2 structural model of the N-terminal region of Scf1 1 reveals a Flo-like fibronectin type III fold characteristic of many eukaryotic adhesion domains. Scf1 is displayed on the surface of S. *cerevisiae* as a fusion with Aga2. (ii). Yeast cells displaying Scf1 are deposited onto coverslips coated with candidate human proteins and spun in a buffered solution. Shear forces detach weakly bound cells. (iii). The resulting radial shear gradient causes cells at the high-shear periphery to detach while cells near the center remain adherent, producing a radial detachment profile which is fit to a sigmoidal equation to derive τ50 **B. Library construction**. The N-terminal Scf1 gene is amplified using mutagenic primers to generate a library of sequence variants. Mutated amplicons are cloned and transformed into *S. cerevisiae* to generate a pooled library of yeast-displayed Scf1 variants. **C. Binding and sorting**. The library is incubated with ligands, followed by incubation with a fluorescent antibody against the ligand. The shift in binding cell population is detected by fluorescence, and FACS separates the cells into four bins ranging from low to high ligand-binding. **D. Epitope identification.** DNA recovered from each sorted bin is analyzed by Illumina sequencing to calculate residue-level fitness scores. Mutational effects are visualized as a heatmap and mapped onto the Scf1 structure to identify ligand-binding epitopes.

Many characterized adhesins engage host substrates through defined ligand-recognition motifs involving specific binding domains or pockets. ^24–26^ In several adhesin systems, including *Candida* species, the adhesin-ligand binding strength increases with applied force, exhibiting so-called catch-bond behavior. This force-activated binding mechanism allows the pathogen to remain attached to host surfaces under physiological shear generated by blood, urine, lymph, airway, or other flow fields, ^27–35^ while allowing the pathogens to release the substrate and remain mobile when flows subside. Here we were interested in studying which ligands are recognized by Scf1 and if these interactions exhibit mechanically stable behavior. Although mutating specific charged and aromatic residues in the N-terminal domain of Scf1 was shown to decrease *C. auris* adhesion to plastics and lipid microparticles, ^19^ the identities of Scf1’s host-associated protein binding partners are not known. Furthermore, it is unclear if these interactions are governed primarily by electrostatics (similarly to their adherence to abiotic surfaces), or if they involve more classical patterns of molecular recognition involving a combination of enthalpic and entropic contributions to binding energy. Given the Flo-like architecture of Scf1 and the prevalence of glycoprotein-rich extracellular matrix (ECM) components at epithelial and vascular surfaces colonized by *C. auris*, we reasoned that human epithelial and ECM proteins represent plausible candidate binding ligands for Scf1.

The N-terminal domain of Scf1 contains regions implicated in adhesion, ^19^ therefore we cloned and displayed this domain on the surface of *S. cerevisiae* yeast (**Fig. 1A** and **1B**) to generate a reductionist platform with which to study biophysical properties and the target of this domain. ^36^ This N-terminal configuration mimics the protein’s orientation on the *C. auris* cell surface. We refer to the yeast-displayed N-terminal domain of full length Scf1 as simply Scf1 in this work. Full sequence information is available in the **Supplementary Information**. This yeast-surface display strategy provided a non-pathogenic model of Scf1-mediated microbial adhesion and enabled expression in a fungal cell-surface context without requiring protein purification. Using quantitative spinning disk adhesion (SDA) assays, ^37–39^ we screened various candidate proteins from human epithelia and ECM tissues for Scf1 binding. This assay allowed us to capture how adhesin-ligand interactions behave under physiological shear that might not be evident from static equilibrium binding assays. We show that yeast-displayed Scf1 binds robustly to human MUC1 and the ECM proteins VIT and FN, but exhibits no detectable binding to collagen I, collagen IV, or fibrinogen.

Interestingly, Scf1 exhibited different ligand binding preferences under static and flow conditions, favoring VIT at equilibrium (i.e. no force) but MUC1 under application of shear stress. The robust binding to human proteins under shear flow is consistent with catch bond-like behavior, wherein applied force stabilizes rather than disrupts adhesive interactions. Finally, to identify the epitope on Scf1 involved in ligand recognition, we adapted a deep mutational scanning (DMS)-based epitope mapping workflow ^40–42^ to map the Scf1 residues involved in MUC1, VIT and FN binding. Binding fitness score heatmaps generated from the DMS datasets revealed shared mutation-sensitive patches for all three ligands, including regions that were previously implicated in adhesion of *C. auris* to abiotic surfaces. ^19^ Analysis of selected single mutants in conventional binding assays validated the DMS-derived epitope and showed reduced binding to all three ligands. A schematic of this workflow is depicted in **Fig. 1**. Together these data provide a structural rationale for broad-spectrum adhesion of Scf1. Overall, these findings identify previously unknown human protein-ligands for Scf1, delineate binding epitopes and establish Scf1 as a broad adhesion regulator that contributes to *C. auris* tissue attachment and colonization in the context of human infections.

## Results

### Scf1 binds to MUC1 and extracellular matrix proteins VIT and FN

Our lab has extensively studied shear-mediated adhesion of microparticles and yeast cells bearing recombinant protein receptors and variants thereof using a custom spinning disk adhesion assay (SDA).^39,43–45^ Using this assay we screened yeast-displayed Scf1 against a panel of human proteins to identify candidate ligands of Scf1 and determine how these interactions behave under physiological shear. This format recapitulates the *in vivo* scenario more accurately than simply assessing equilibrium binding under static conditions. The first binding panel included the following human proteins in their native format (i.e. isolated from human tissues): MUC1, VIT, FN, collagen I, collagen IV and fibrinogen. Although non-exhaustive, these ligands represent a diverse set of proteins found at epithelial and tissue interfaces, and are commonly implicated in adhesive colonization by microbes. ^46^

Yeast cells displaying Scf1 were seeded onto circular cover glasses coated with the target proteins (**Fig. 1A**). We subjected the samples to a controlled spinning procedure using a custom-built device, which generates shear flow that increases radially outward from the center point on the disk to simulate physiological fluid flow. Under the conditions we report, this assay applies shear stresses from 0 to 500 dyn/cm², covering the physiological range (0.2–120 dyn/cm²) and higher. Based on the fluid dynamics of the system the shear stress is highest at the periphery of the cover glass and lowest at the center. This results in a radial distribution of cell density with high cell density at the center and low density towards the edges. Following spinning, the cover glasses were raster imaged under a microscope and the images were processed using a computational image analysis pipeline to generate cell-density distribution heatmaps that visualize the radial detachment pattern (**Fig. 1A**).

To test which members of a human protein panel are recognized by Scf1, we modified cover glasses with recombinant MUC1, VIT, FN, collagen I, collagen IV, and fibrinogen using epoxysilane chemistry. After allowing Scf1-bearing yeast cells to adhere to the surfaces, they were exposed to a shear stress spinning protocol (3000 rpm, 5 minutes) and followed by imaging and analysis. As shown in **Fig 2A**, the cover glasses coated with MUC1, VIT and FN exhibited clear ligand-dependent adhesion, with adherent cells decreasing radially from the low-shear center to the high-shear periphery. This effect was most prominent for MUC1, followed by VIT and FN and was consistent across multiple independent experiments. In contrast, heatmaps for collagen 1, IV and fibrinogen exhibited low and patchy cell attachment across the cover glasses, indicating weak or no binding under our experimental conditions **(Fig. 2A)**.

**Figure 2.**
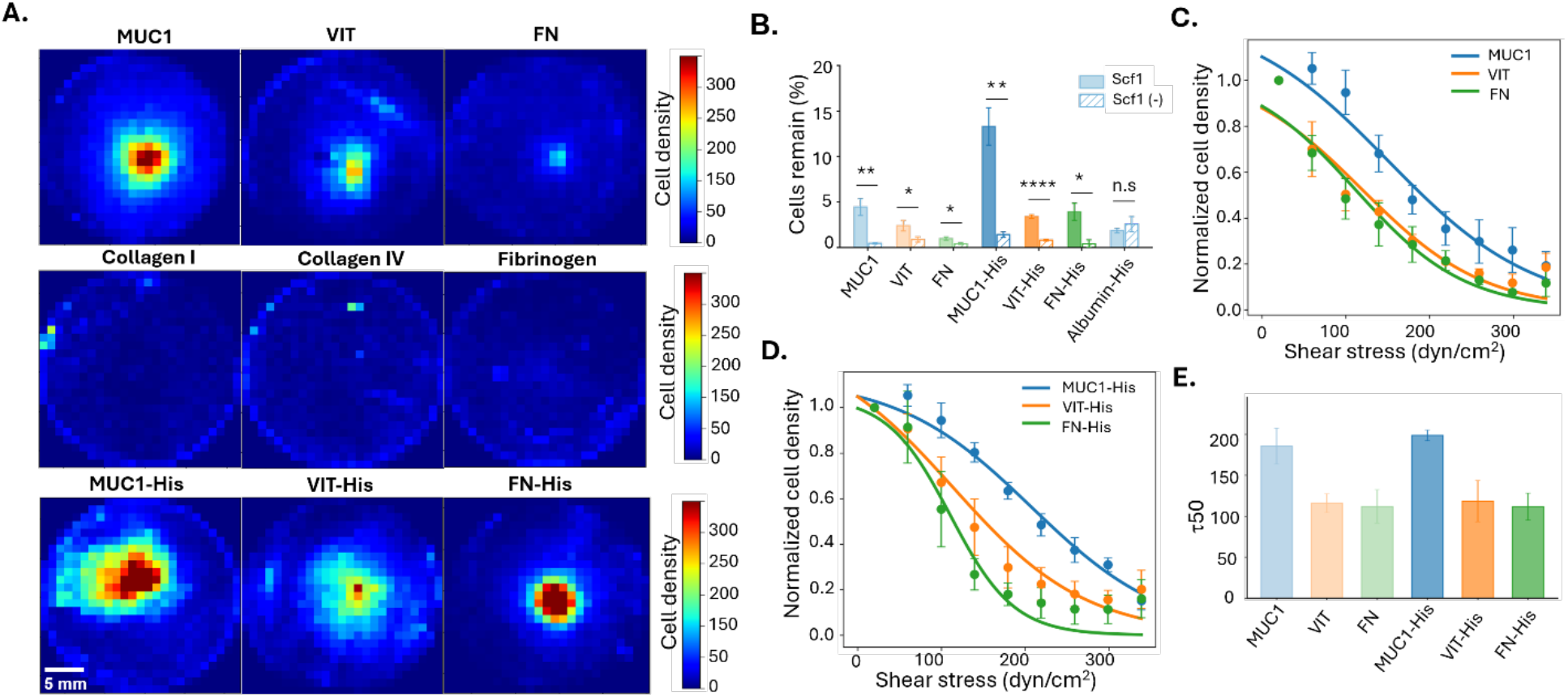
Scf1 binds to ECM ligands under shear stress. **A**. **Cell-density distribution heatmaps**. Heatmaps show the spatial density of yeast cells displaying Scf1 that remained attached to 25-mm cover glasses coated with different ECM ligands after spinning (3000 rpm, 5 min.). For visualization, each post-spin image was partitioned into 25 × 25–pixel bins. Cells in each bin were counted and mapped to a colormap. Color scale shows number of detected cells per spatial bin with warmer colors indicating higher local cell density (0–300 cells/bin; 5 mm scale bar in the lower left). The rotation axis is typically at the center of each panel, and shear stress increases with radius. Panels show superimposed (averaged) heatmaps from three to four independent experiments for individual ECM ligands: MUC1, VIT, FN, collagen IV, fibrinogen, MUC1 (HIS), VIT (HIS), and FN (HIS). **B**. **Percentage of cells remaining after spinning**. Bar plots showing comparison of percentage of cells that remain adhered after spinning cover glasses between Scf1 and Scf1(-) yeasts. **C**. **Cell density versus shear stress for native proteins**. Normalized cell densities of yeast cells displaying Scf1 (from **A**) are plotted against shear stress (dyn/cm^2^). Data are fit to a sigmoidal function to quantify the adhesion strengths of yeast-displayed Scf1 to native MUC1 (blue curve-fit), VIT (orange curve-fit) and FN (green curve-fit). **D**. **Cell density versus shear stress for His-tagged protein**s. Normalized cell densities of yeast cells displaying Scf1 are plotted against shear stress (dyn/cm^2^). MUC1-His (blue curve-fit), VIT-His (orange curve-fit) and FN-His (green curve-fit). **E. Quantification of adhesion strengths**. Adhesion strengths of Scf1 for each ligand expressed as the average τ₅₀ value. Error bars in panels B, C, D and E represent SEM from four to five independent experiments.

To further validate that the observed adhesion was due to specific Scf1-ligand interactions, we tested the binding of yeast-displayed Scf1 against a panel of recombinantly expressed and purified proteins. This panel included commercially available forms of MUC1, VIT, FN and albumin, each carrying a C-terminal His-tag, expressed and purified in HEK 293 cells. Consistent with the results from the human tissue-sourced native ligands, the three His-tagged ligands reproduced the same adhesion strength hierarchy (MUC1 > VIT > FN). To further validate specificity of the interactions, as a negative control, we tested yeasts that displayed cohesin protein domain of CipA (from *Clostridium thermocellum*), an unrelated folded protein with no known mammalian ECM-binding activity, in place of Scf1. This construct, referred to as Scf1(-), used the same vector backbone, HA-tag and Aga2 protein. This design allowed us to evaluate whether differences in adhesion were due to the displayed Scf1 rather than the Aga2p display scaffold alone. For each ligand, we compared the percentage cells remaining adhered after spinning between Scf1-displaying yeasts and Scf1(-) yeasts on the same coated surface (**Fig. 2B, Fig. S1**). The Scf1(-) yeasts showed significantly weaker adhesion for all three binding ligands (MUC1 (*p* = 0.0034) VIT (*p* = 0.0399), FN (*p* = 0.0210), MUC1-His (*p* = 0.0020), VIT-His (*p* = 5.8E-06) and FN-His (*p* = 0.0483)). In contrast, the Scf1 versus Scf1(-) comparison was not significant (n.s.) for albumin-His (*p* = 0.318) suggesting that Scf1 adhesion to albumin adhesion was non-specific under these conditions (**Fig. 2B, Fig. S2**).

We next fitted the cell-density distribution data from **Fig. 2A** to a sigmoid model, allowing us to derive the *τ50* parameter, which represents the shear stress at which 50% of the cells remain adherent. This allowed us to quantify the adhesion strength of yeast-displayed Scf1 to MUC1, VIT and FN (**Fig. 2C-D**). For the native proteins, yeast-displayed Scf1 exhibited the greatest adhesion strength on MUC1 (τ50 = 186 ± 22 dyn/cm^2^), followed by VIT (τ50 = 116 ± 11 dyn/cm^2^) and FN (τ50 = 112 ± 20 dyn/cm^2^) (**Fig. 2C** and **2D**). Similarly, the recombinant His-tagged ligands reproduced the same adhesion strength hierarchy; MUC1-His showed the highest adhesion strength (τ50 = 198 ± 7 dyn/cm^2^), followed by VIT-His (τ50 = 118 ± 25 dyn/cm^2^) and FN-His (τ50 = 112 ± 16 dyn/cm^2^). Statistical analysis showed that while τ50 values for MUC1 do not significantly differ from FN (*p* = 0.08), it was significantly higher than VIT (*p* = 0.04). For the His-tagged ligands, however, the τ50 values for MUC1-His were significantly higher than both VIT-His (*p* = 0.02) and FN-His (*p* = 0.008). Overall, these data show that Scf1 binds MUC1-containing surfaces under shear stress, with lesser adhesion also found for VIT and FN.

The His-tagged proteins also allowed us to validate the results from the surface-based adhesion assays via an in-solution, FACS-based binding assay (**Fig. 3A**). Here, yeasts displaying Scf1 were titrated with increasing concentrations of same set of His-tagged proteins used for the spinning-disk assay (MUC1-His, VIT-His and FN-His), and bound ligand was detected via anti-HIS primary antibody and a fluorescent secondary antibody. For all three ligands, we observed a ligand-dependent increase in median fluorescence consistent with a specific Scf1-ligand interaction. Albumin-His served as a negative control and showed a negligible increase in fluorescence comparable to that observed for Scf1(-) yeasts (**Fig. 3B**). It is notable that the hierarchy of ligand binding strength observed under shear (**Fig. 2**) was not recapitulated in the equilibrium binding assays. Yeast-displayed Scf1 binds to VIT-His with the highest apparent binding affinity (*K_D_* _app_ 14 ± 3 nM), followed by FN-His (*K_D_*_app_ 75 ± 9 nM), whereas MUC1-His (*K_D_*_app_ 215 ± 16 nM) showed the weakest binding. Additionally, VIT-His showed the largest increase in median fluorescence. In summary, the spinning-disk and FACS-based assays capture different aspects of the Scf1-ligand binding behavior, with surface-immobilized measurements under shear flow likely reflecting more physiologically relevant interactions.

**Figure 3.**
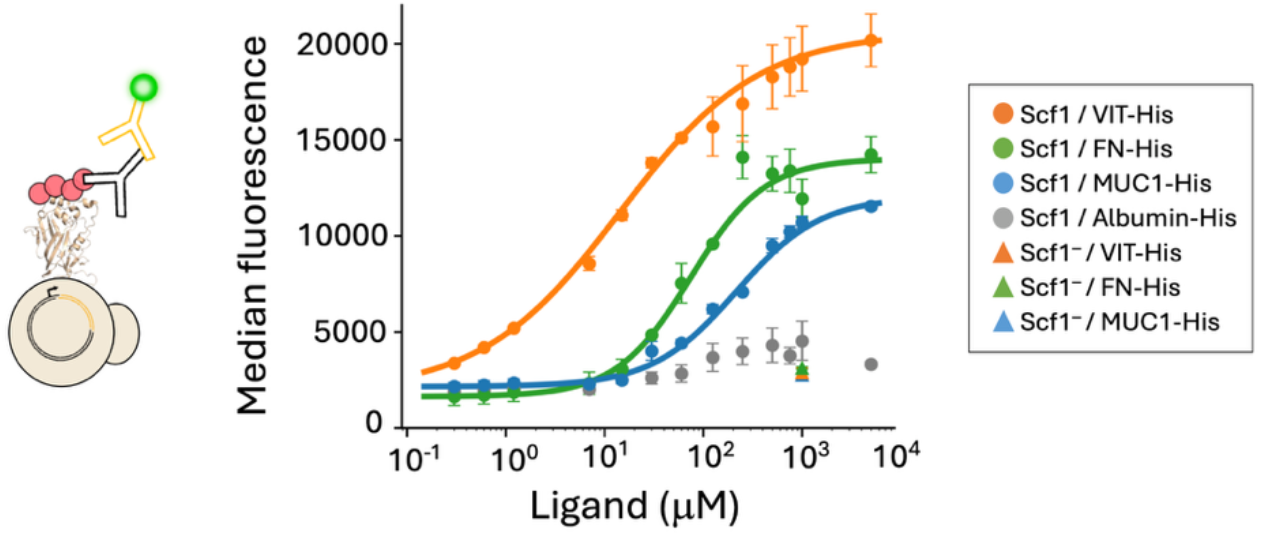
Scf1 binds to host-associated proteins at equilibrium. The schematic of FACS-based binding assay is shown as an inset. Yeast-displaying Scf1 were incubated with His-tagged ligands. Binding of Scf1 to the His-tagged ligands was detected by flow cytometry using fluorescent anti-His antibodies. Plot shows median fluorescence intensity measured by flow cytometry for yeast displaying Scf1, incubated with increasing concentrations of MUC1-His, VIT-His, FN-His, and albumin-His. Binding signal from 1 μM of each ligand for Scf1(-) is shown as triangles. Data points were fitted to the Hill equation using a custom python script. Error bars indicate SEM from 3–4 independent experiments.

### DMS-based epitope mapping identifies shared recognition sites for MUC1, VIT, and FN

We next sought to identify the residues within the N-terminal domain responsible for ligand recognition. To this end, we performed DMS on yeast-displayed Scf1 and used a FACS sorting and DNA sequencing pipeline that was modified and adapted from prior work in our laboratory ^40,42^ (**Fig. 1, panels B–D**). In brief, we constructed a near-comprehensive single-substitution variant library spanning the Scf1 adhesive domain (residues 28–255) using the one-pot mutagenesis method ^47^ with NNN mutagenic primers and displayed the library on the surface of *S. cerevisiae*. Following the experimental rationale shown in **Fig. 3**, we incubated the library with each ligand (His-tagged form) slightly above the K_D_. The shift in cell population was detected by antibody immunostaining against the His-tag and cells were sorted using FACS into four bins spanning low to high fluorescent binding signals. The variant composition of each bin was then determined by Illumina sequencing, and a per-residue fitness score was computed for every position (see Methods) according to previously established protocols. ^40^ Higher fitness values reflected enrichment of substitutions that retain (or enhance) binding and lower fitness reflects depletion of binding-competent variants.

The displayed construct comprises residues 28–255 of full-length Scf1, which we renumbered as positions 1–228 for DMS analysis. The residue numbers reported here therefore refer to construct positions in our gene cassette, unless otherwise specified. Comparison of the binding fitness maps (normalized to expression) for MUC1, VIT, and FN revealed several recurrent clusters of binding-sensitive substitutions (**Fig. 4A**) in Scf1. Upon visual inspection, the most prominent region sensitive to mutations for all three ligands comprised a cation-aromatic patch of residues from 120 to 125 (residues 147-152 of native Scf1; AAs. KRHKHK), where multiple substitutions reduced binding to all three ligands. Additional shared mutation-sensitive patches were observed at residues 14–18 (residues 41-45 of native Scf1; aa. TPSKH), 25–28 (residues 52-55 of native Scf1; aa. HHRR), 135–140 (residues 162-167 of native Scf1; aa. HKLEVR) and 214-216 (residues 241-243 of native Scf1) (**Fig. 4B**).

**Figure 4.**
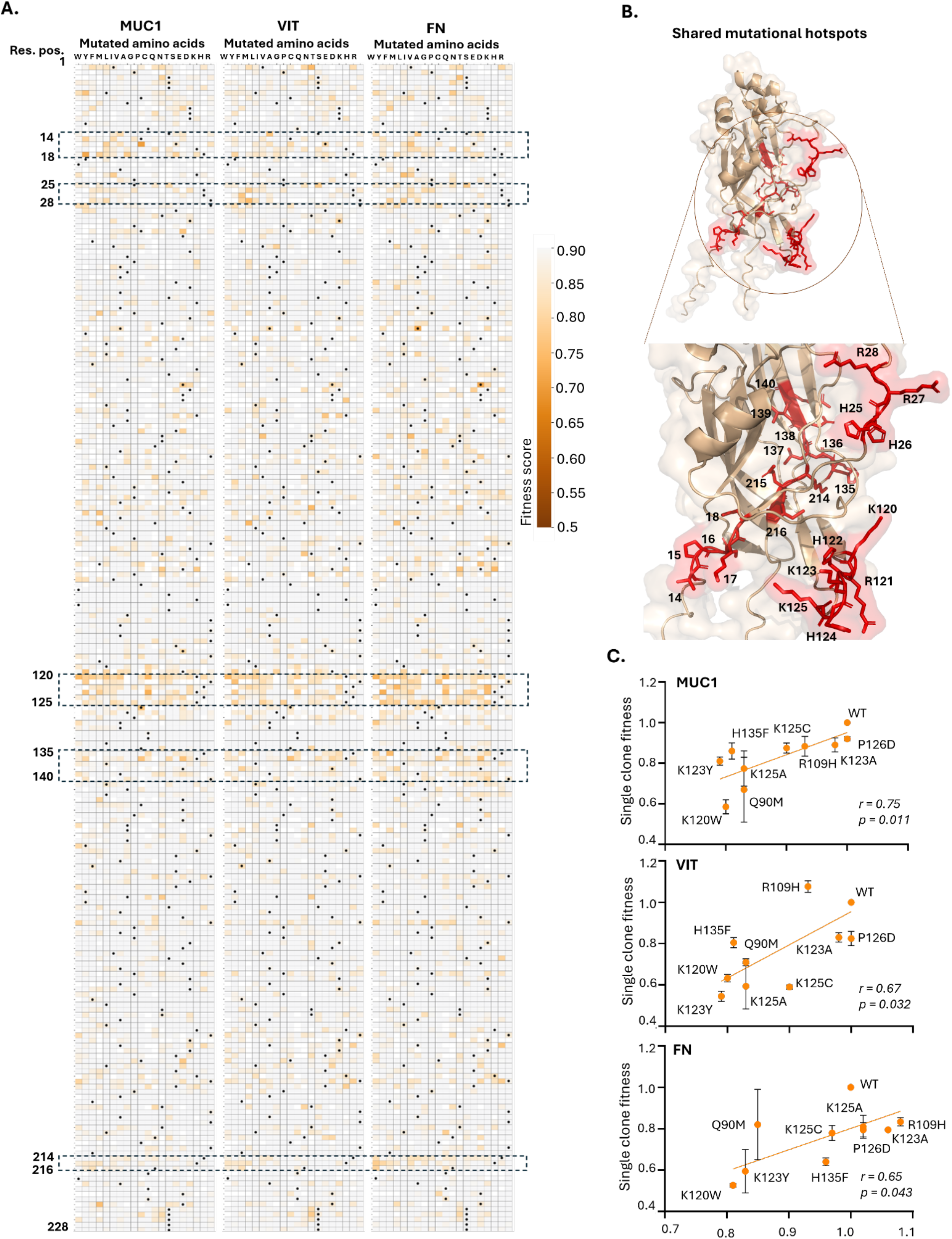
Deep mutational scanning identifies shared ligand-binding epitopes in the Scf1 N-terminal adhesive domain. A. Fitness heatmaps. Expression-normalized DMS fitness maps for Scf1 binding to MUC1 (left), VIT (center), and FN (right). Each row represents a residue position in the yeast-displayed Scf1 construct, and each column represents one of the 20 amino-acid identities, with black dots marking the wild-type Scf1 residue at each position. Orange shading indicates substitutions associated with reduced ligand binding. Dashed boxes highlight recurrent mutation-sensitive patches identified across the three ligands. **B. Shared mutational hotspots**. Mapping of the recurrent mutation-sensitive regions onto the Scf1 AlphaFold2-predicted structure. Residues within the identified patches are shown in red. The enlarged view highlights spatial clustering of residues from multiple discontinuous, surface and core, patches. The prominent 120–125 and 25-28 patches involved in abiotic surface binding form a contiguous surface-accessible interface and are labeled with both residue numbers and amino-acid identities. The remaining mutation-sensitive positions are labeled by residue number only for clarity. **C. Single-clone validation**. Experimental validation of selected mutations identified by DMS. Individually reconstructed Scf1 variants were assessed for binding to MUC1, VIT, and FN, and their measured binding-fitness were correlated with the corresponding DMS-derived fitness scores to derive the pearson correlation coefficients (MUC1, *r* = 0.75; VIT, *r* = 0.67; FN, *r* = 0.65). The correlation between the DMS and single-clone fitness was statistically significant (MUC1, *p* = 0.011; VIT, *p* = 0.032; FN, *p* = 0.043)

To analyze these putative binding patches in detail, we calculated a position-level median fitness change. At each residue position, the median fitness was calculated across all measured non-synonymous substitutions, and the fitness change was defined as 1 − median(fitness), where a fitness score of 1 represents WT-like binding. We then calculated the patch-level value by averaging these position-level fitness changes across all residues within the predefined patch. Thus, a positive patch-level value indicates that the median fitness across substitutions was, on average, below WT at positions within that patch. Second, we calculated the fraction of substitutions with fitness below WT at each position as the number of measured non-synonymous substitutions with fitness <1 divided by the total number of measured non-synonymous substitutions. The patch-level fraction below WT was then calculated by averaging these position-level fractions across all evaluable residues within each predefined patch, for example patches 14–18 and 120–125. For comparison, non-patch background values for both metrics were calculated in the same manner across all evaluable residues outside the predefined patches.

This analysis revealed that patch 120-125 is the strongest shared mutation-sensitive patch by both metrics. Depending on the ligand, the average position-level median fitness reduction across this patch was 6–9%: 9% for MUC1, 6% for VIT, and 7% for FN. Independently, the mean position-level fraction of substitutions with fitness below WT was 79–86%: 86% for MUC1, 81% for VIT, and 79% for FN, compared with 43– 48% in the non-patch background. Thus, the 79–86% values describe the proportion of substitutions that reduced fitness, not the magnitude of the reduction in binding. The other four mutation-sensitive patches showed smaller median fitness reductions, generally 1–3%, but still exhibited higher fractions of below-WT substitutions, ranging from 61% to 75%, than the non-patch background (**Table 1**). Interestingly, the residues R27–R28 (of patch 25-28) corresponding to R54–R55 in native Scf1, were previously implicated in Scf1-mediated adhesion to plastics.^19^ Structurally, both the prominent 120–125 and the 25-28 are neighboring mutation-sensitive regions, forming a contiguous surface-accessible interface, whereas the other three patches comprise the core of the domain (**Fig. 4B**). For the non-patch background, the average of the position-level median fitness scores was slightly above 1: 1.0096 for MUC1, 1.0171 for VIT, and 1.0179 for FN. Consequently, the corresponding background values for the median-fitness-change metric were slightly negative: approximately −1%, −2%, and −2%, respectively. These small deviations are best interpreted as essentially WT-like background fitness rather than biologically meaningful increases in binding. Using the second metric, for the non-patch background, the mean position-level fraction of non-synonymous substitutions with fitness below 1 was 48% for MUC1, 43% for VIT, and 44% for FN.

**Table 1:**
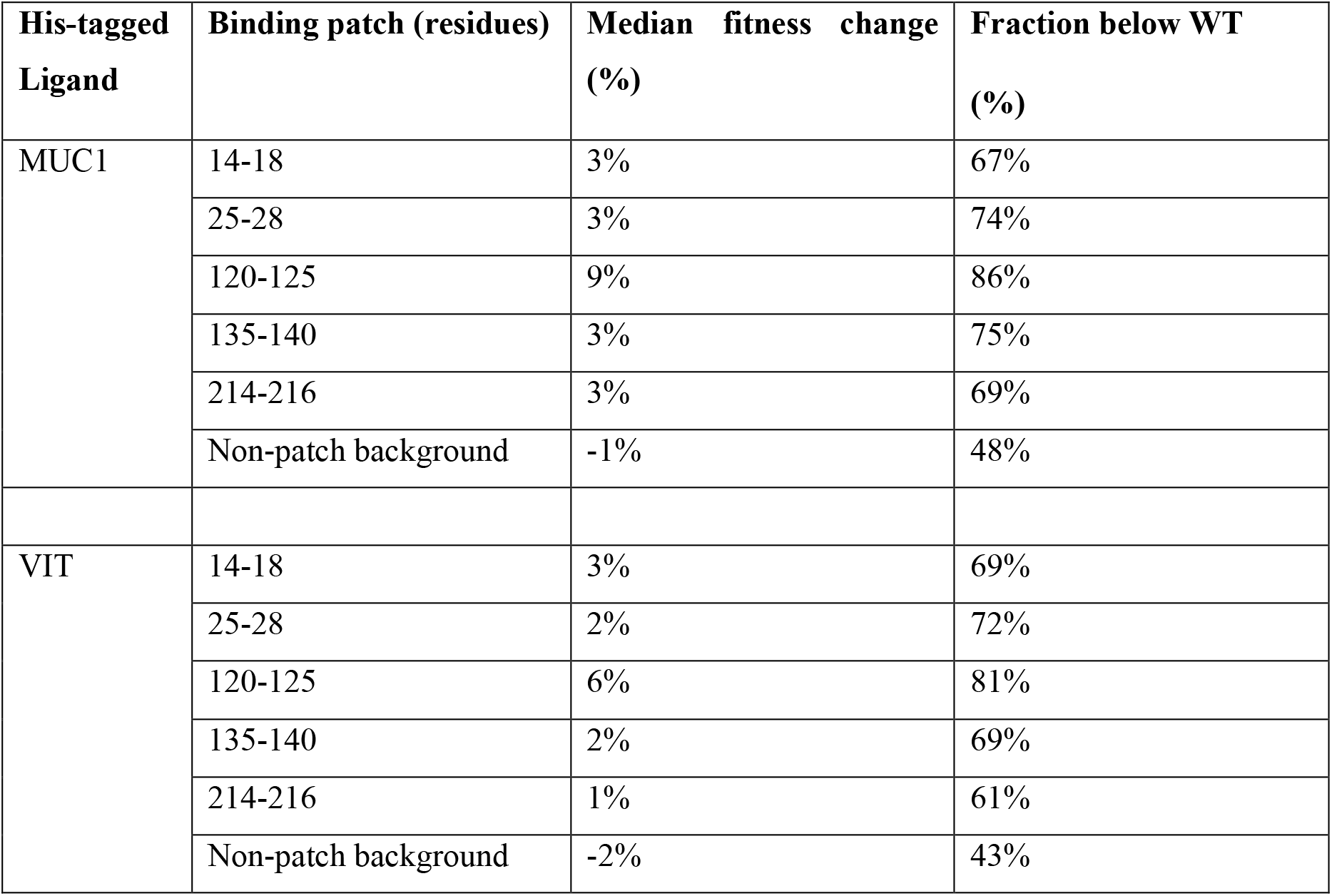
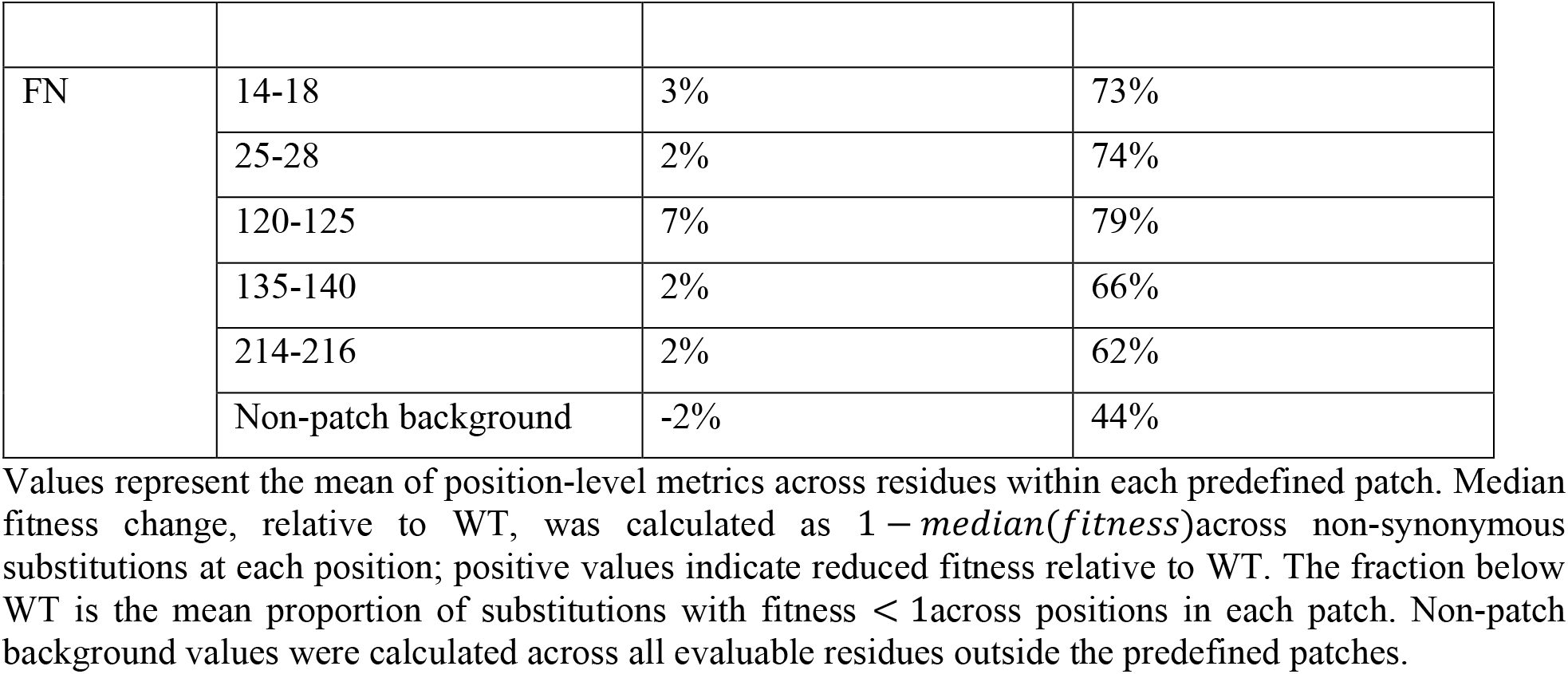
Patch-level median fitness reduction and fraction of activity-reducing substitutions.

| **His-tagged**<br>**Ligand** | **Binding patch (residues)** | **Median fitness change**<br>**(%)** | **Fraction below WT**<br>**(%)** |
| --- | --- | --- | --- |
| MUC1 | 14-18 | 3% | 67% |
|  | 25-28 | 3% | 74% |
|  | 120-125 | 9% | 86% |
|  | 135-140 | 3% | 75% |
|  | 214-216 | 3% | 69% |
|  | Non-patch background | -1% | 48% |
| VIT | 14-18 | 3% | 69% |
|  | 25-28 | 2% | 72% |
|  | 120-125 | 6% | 81% |
|  | 135-140 | 2% | 69% |
|  | 214-216 | 1% | 61% |
|  | Non-patch background | -2% | 43% |
| FN | 14-18 | 3% | 73% |
|  | 25-28 | 2% | 74% |
|  | 120-125 | 7% | 79% |
|  | 135-140 | 2% | 66% |
|  | 214-216 | 2% | 62% |
|  | Non-patch background | -2% | 44% |

Values represent the mean of position-level metrics across residues within each predefined patch. Median fitness change, relative to WT, was calculated as $1 - \text{median}(\text{fitness})$ across non-synonymous substitutions at each position; positive values indicate reduced fitness relative to WT. The fraction below WT is the mean proportion of substitutions with fitness $< 1$ across positions in each patch. Non-patch background values were calculated across all evaluable residues outside the predefined patches.

To validate the pooled DMS measurements, we individually reconstructed ten variants containing mutations from the identified epitope regions, most from the 120–125 binding patch, and assessed ligand binding using monoclonal yeast cultures. The single-clone fitness correlated significantly with DMS fitness for all three ligands MUC1 (Pearson r = 0.75; p (two-tailed) = 0.0117, VIT (Pearson r = 0.68; p (two-tailed) = 0.0323) and FN (Pearson r = 0.65; p (two-tailed) = 0.0438). Overall, most mutants tested showed reduced fitness across all three ligands, suggesting a substantial overlap in their functional recognition surfaces. However, we also observed differences in the magnitude of individual mutational effects suggesting some ligand-dependent contributions within these shared interfaces. These results suggest a role for the identified residues in ligand recognition rather than a general loss of protein folding or surface display (**Supplementary Information**), although effects on local structure cannot be excluded. Together, these findings identify shared regions that contribute to Scf1 recognition of MUC1, VIT, and FN, with the main binding patch being the surface-exposed, flexible loop composed of cationic and aromatic residues (KRHKHK).

## Discussion

In this work, by combining SDA assays with FACS-based binding measurements, we demonstrate that the N-terminal domain of Scf1 binds to a structurally diverse set of proteins including MUC1, VIT, and FN. Under shear stress, we found that Scf1 exhibits the strongest interaction with MUC1 as compared to VIT and FN, while no appreciable interactions were detected for collagen I, IV, fibrinogen, and albumin. Our binding experiments involving yeast cells displaying Scf1 on the cell surface mimic the natural presentation of the Scf1 adhesin on *C. auris* in a format that is experimentally tractable for biophysical assays and DMS. The binding behavior that we measured reflects the multivalent nature of the binding interactions that occur during host colonization by cell-surface adhesins. DMS further identified shared epitopes on the surface of the Scf1 N-terminal domain involved in recognition of the three ligands, providing a molecular basis for understanding how Scf1 combines broad ligand recognition with substrate- and force-dependent adhesion.

### MUC1-dominant binding with complementary binding to VIT and FN

Our decision to test MUC1 as a potential binder was guided by the observation that Scf1 adhesin demonstrates structural parallels to *Vibrio cholerae* adhesion systems in which mucin-binding (often glycan-recognizing) modules promote persistence at host barriers. ^48,49^ In some instances, mucins and mucin-derived glycans can also act as decoy receptors, attenuating fungal pathogenic traits in *C. albicans* and suppressing virulence phenotypes of adhesion and biofilm formation. ^50–52^ Therefore, the strong interaction of Scf1 with MUC1 under shear stress suggests that *C. auris* may target mucin-rich niches such as skin or mucosal surfaces for initial adhesion. The finding that Scf1 binds to FN and VIT suggests a potential role in *C. auris* attachment to abiotic materials, such as indwelling medical-device coated with VIT and FN adsorbed from surrounding body fluids, ^53–55^ and wound or tissue-remodeling environments, where provisional matrix contains these proteins. ^56^ In terms of the non-binding ligands, we found that yeast-displayed Scf1 demonstrates low retention on the cover-glasses coated with fibrinogen, collagen I, IV, and albumin. This does not entirely exclude weak binding to these ligands in other contexts, but adhesion to these ligands was not observed under the shear-based assay conditions tested here. Further studies should examine additional ligands as potential host substrates across diverse biological contexts.

### Electrostatic promiscuity without loss of mechanical specificity: Toward a cationic catch-bond paradigm

Given that Scf1 governs adhesion using primarily electrostatic interactions, ^19^ it is plausible that Scf1 interacts with charged patches, such as acidic residues of host-associated proteins and/or negatively charged sialylated or sulfated glycans on protein surfaces. However, the ligand-specific differences observed across structurally unrelated host proteins and the altered ligand preference under shear suggest that simple electrostatic attraction alone may not fully explain Scf1 adhesion behavior. Further, the rank-order differences between adhesion strength under shear stress vs. binding at equilibrium suggest Scf1 interacts with distinct ligands in ways that are not captured by a single measure of binding strength. For example, VIT may support stronger apparent binding to cell-surface-displayed Scf1 when the ligand is free in solution under equilibrium conditions, whereas MUC1 may form interactions with Scf1 that persist more effectively under shear stress. Taken together, these findings extend the current model of Scf1-mediated adhesion beyond electrostatic surface association to abiotic surfaces by showing the Scf1 interacts with host-ligands in a ligand- and force-dependent manner.

### Shared recognition regions may support adhesion across diverse abiotic and biotic environments

The DMS data suggests that a shared set of functional regions are crucial for Scf1-mediated binding to the structurally diverse human proteins MUC1, VIT and FN. Validation of DMS data by single clone analysis showed that most of the mutations tested reduced binding activity across all three ligands. However, the magnitude of the individual mutational effects varied between MUC1, VIT, and FN. This suggests that residues within this shared binding epitope contribute differently to interactions with the individual ligands (**Fig. 4C**). The overlap between the binding residues identified by DMS (patch 25-28, **Fig. 4B**) and the arginine pair previously reported by Santana et al. ^19^ as being required for *C. auris* adhesion to plastic surfaces has two-fold significance. First, it suggests that Scf1 uses overlapping epitopes to interact with both human proteins and abiotic materials, rather than relying on separate mechanisms. Such versatility could be advantageous for *C. auris*, which must transition between medical devices, hospital surfaces, skin, mucosal or tissue-associated environments. Second, this provides independent support that our yeast display/DMS workflow can detect determinants of pathogenic adhesion that are relevant in native physiological contexts. More broadly, the presence of an overlapping functional interface raises the possibility that targeting a limited region of Scf1 could inhibit adhesion to multiple host substrates and potentially reduce both host colonization and attachment to medically relevant surfaces. Taken together, these results identify a promiscuous adhesive epitope in which several discontinuous patches contribute to Scf1–ligand binding and mechanical stabilization.

## Materials and Methods

The N-terminal adhesive domain of Scf1 (residues 28–255) was displayed on the surface of *Saccharomyces cerevisiae* and used to characterize ligand recognition by complementary cell-based and sequencing approaches. Adhesion to candidate human proteins was quantified under hydrodynamic shear using a spinning-disk adhesion assay, while equilibrium binding to His-tagged ligands was measured by flow cytometry. To identify residues contributing to ligand recognition, we generated a deep mutational scanning library of Scf1, linked individual variants to DNA barcodes using PacBio long-read sequencing, and quantified expression and ligand-binding fitness by FACS sorting followed by Illumina sequencing. Selected DMS variants were reconstructed individually for validation, and mutation-sensitive regions were mapped onto the predicted Scf1 structure. Full experimental procedures, data-processing and statistical methods are provided in the supplementary section.

## Supplementary information

Supplementary information contains detailed description of methods, flow-cytometric validation of Scf1 and Scf1(−) surface expression (**Fig. S1**), spinning-disk adhesion controls comparing Scf1 and Scf1(−) (**Fig. S2**), representative DMS expression and ligand-binding sorting profiles (**Fig. S3**), replicate reproducibility and fitness-score distributions for the DMS datasets (**Fig. S4**), surface-expression analysis of individually reconstructed DMS variants (**Fig. S5**), and the complete gene and protein sequences of the yeast-display constructs (**Tables S1 and S2**).

## Acknowledgments

This work was supported by the University of Basel, ETH Zurich and a Consolidator Grant (MB22.00050) from the Swiss State Secretariat for Education, Research and Innovation (SERI) to MN.

## References

1. Manolakaki, D. et al. Candida infection and colonization among trauma patients. Virulence 1, 367– 375 (2010).

2. Brandt, M. E. & Lockhart, S. R. Recent Taxonomic Developments with Candida and Other Opportunistic Yeasts. Curr. Fungal Infect. Rep. 6, 170–177 (2012).

3. Chowdhary, A., Jain, K. & Chauhan, N. Candida auris Genetics and Emergence. Annu. Rev. Microbiol. 77, 583–602 (2023).

4. Schelenz, S. et al. First hospital outbreak of the globally emerging Candida auris in a European hospital. Antimicrob. Resist. Infect. Control 5, 35–35 (2016).

5. Lockhart, S. R. et al. Simultaneous Emergence of Multidrug-Resistant Candida auris on 3 Continents Confirmed by Whole-Genome Sequencing and Epidemiological Analyses. Clin. Infect. Dis. 64, 134– 140 (2017).

6. Satoh, K. et al. Candida auris sp. nov., a novel ascomycetous yeast isolated from the external ear canal of an inpatient in a Japanese hospital. Microbiol. Immunol. 53, 41–44 (2009).

7. Bing, J. et al. Global emergence and rapid spread of Candidozyma auris (syn. Candida auris): epidemiology, biology, and antifungal resistance. Clin. Microbiol. Rev. e0039425 (2026).

8. CDC. Tracking C. auris. Candida auris (C. auris) https://www.cdc.gov/candida-auris/tracking-c-auris/index.html (2026).

9. Sabino, R., Veríssimo, C., Pereira, Á. A. & Antunes, F. Candida auris, an Agent of Hospital-Associated Outbreaks: Which Challenging Issues Do We Need to Have in Mind? Microorganisms 8, (2020).

10. Smoak, R. A., Snyder, L. F., Fassler, J. S. & He, B. Z. Parallel expansion and divergence of an adhesin family in pathogenic yeasts. Genetics 223, 1–19 (2023).

11. Welsh, R. M. et al. Survival, Persistence, and Isolation of the Emerging Multidrug-Resistant Pathogenic Yeast Candida auris on a Plastic Health Care Surface. J. Clin. Microbiol. 55, 2996–3005 (2017).

12. Li, J., Brandalise, D., Coste, A. T., Sanglard, D. & Lamoth, F. Exploration of novel mechanisms of azole resistance in Candida auris. Antimicrob. Agents Chemother. 68, e0126524 (2024).

13. Ware, A. et al. Dry Surface Biofilm Formation by Candida auris Facilitates Persistence and Tolerance to Sodium Hypochlorite. APMIS 133, (2025).

14. Horton, M. V. & Nett, J. E. Candida auris Infection and Biofilm Formation: Going Beyond the Surface. Curr. Clin. Microbiol. Rep. 7, 51–56 (2020).

15. Kappel, D. et al. Genomic epidemiology describes introduction and outbreaks of antifungal drug-resistant Candida auris. NPJ Antimicrob. Resist. 2, 26 (2024).

16. Louvet, M. et al. Ume6-dependent pathways of morphogenesis and biofilm formation in Candida auris. Microbiol. Spectr. 12, e0153124 (2024).

17. de Groot, P. W. J., Bader, O., de Boer, A. D., Weig, M. & Chauhan, N. Adhesins in human fungal pathogens: glue with plenty of stick. Eukaryot. Cell 12, 470–481 (2013).

18. Datta, A. et al. Differential skin immune responses in mice intradermally infected with Candida auris and Candida albicans. Microbiol. Spectr. 11, e02215–23 (2023).

19. Santana, D. J. et al. A Candida auris–specific adhesin, Scf1, governs surface association, colonization, and virulence. Science 381, 1461–1467 (2023).

20. Wang, T. W. et al. Functional redundancy in Candida auris cell surface adhesins crucial for cell-cell interaction and aggregation. Nat. Commun. 15, 9212–9212 (2024).

21. Kimizuka, F. et al. Role of type III homology repeats in cell adhesive function within the cell-binding domain of fibronectin. J. Biol. Chem. 266, 3045–3051 (1991).

22. Bencharit, S. et al. Structural insights into fibronectin type III domain-mediated signaling. J. Mol. Biol. 367, 303–309 (2007).

23. Kinoshita, T. Glycosylphosphatidylinositol (GPI) Anchors: Biochemistry and Cell Biology: Introduction to a Thematic Review Series. Journal of lipid research vol. 57 4–5 Preprint at 10.1194/jlr.E065417 (2016).

24. Salgado, P. S. et al. Structural basis for the broad specificity to host-cell ligands by the pathogenic fungus Candida albicans. Proc. Natl. Acad. Sci. U. S. A. 108, 15775–15779 (2011).

25. Foster, T. J. The MSCRAMM family of cell-wall-anchored surface proteins of gram-positive cocci. Trends Microbiol. 27, 927–941 (2019).

26. Hoffmann, D. et al. Functional reprogramming of Candida glabrata epithelial adhesins: the role of conserved and variable structural motifs in ligand binding. J. Biol. Chem. 295, 12512–12524 (2020).

27. Chan, C. X. J. & Lipke, P. N. Role of force-sensitive amyloid-like interactions in fungal catch bonding and biofilms. Eukaryot. Cell 13, 1136–1142 (2014).

28. Zhu, C., Lou, J. & McEver, R. P. Catch bonds: physical models, structural bases, biological function and rheological relevance. Biorheology 42, 443–462 (2005).

29. Sokurenko, E. V., Vogel, V. & Thomas, W. E. Catch-bond mechanism of force-enhanced adhesion: counterintuitive, elusive, but … widespread? Cell Host Microbe 4, 314–323 (2008).

30. Kim, J., Zhang, C.-Z., Zhang, X. & Springer, T. A. A mechanically stabilized receptor-ligand flex-bond important in the vasculature. Nature 466, 992–995 (2010).

31. Milles, L. F., Schulten, K., Gaub, H. E. & Bernardi, R. C. Molecular mechanism of extreme mechanostability in a pathogen adhesin. Science 359, 1527–1533 (2018).

32. Liu, Z. et al. High force catch bond mechanism of bacterial adhesion in the human gut. Nat. Commun. 11, 4321–4321 (2020).

33. Mathelié-Guinlet, M., Viela, F., Alsteens, D. & Dufrêne, Y. F. Stress-induced catch-bonds to enhance bacterial adhesion. Trends Microbiol. 29, 286–288 (2021).

34. Alsteens, D., Garcia, M. C., Lipke, P. N. & Dufrêne, Y. F. Force-induced formation and propagation of adhesion nanodomains in living fungal cells. Proc. Natl. Acad. Sci. U. S. A. 107, 20744–20749 (2010).

35. Liu, H., Liu, Z., Yang, B., Lopez Morales, J. & Nash, M. A. Optimal Sacrificial Domains in Mechanical Polyproteins: S. epidermidis Adhesins Are Tuned for Work Dissipation. JACS Au 2, 1417–1427 (2022).

36. Teymennet-Ramírez, K. V., Martínez-Morales, F. & Trejo-Hernández, M. R. Yeast Surface Display System: Strategies for Improvement and Biotechnological Applications. Front. Bioeng. Biotechnol. 9, (2022).

37. Mohandas, N., Hochmuth, R. M. & Spaeth, E. E. Adhesion of red cells to foreign surfaces in the presence of flow. J. Biomed. Mater. Res. 8, 119–136 (1974).

38. García, A. J., Ducheyne, P. & Boettiger, D. Quantification of cell adhesion using a spinning disc device and application to surface-reactive materials. Biomaterials 18, 1091–1098 (1997).

39. Santos, M. S., Liu, H., Schittny, V., Vanella, R. & Nash, M. A. Correlating single-molecule rupture mechanics with cell population adhesion by yeast display. Biophys. Rep. (N. Y.) 2, (2022).

40. Vanella, R. et al. Understanding activity-stability tradeoffs in biocatalysts by enzyme proximity sequencing. Nat. Commun. 15, 1807 (2024).

41. Gomes, D. E. B., Yang, B., Vanella, R., Nash, M. A. & Bernardi, R. C. Integrating Dynamic Network Analysis with AI for Enhanced Epitope Prediction in PD-L1:Affibody Interactions. J. Am. Chem. Soc. 146, 23842–23853 (2024).

42. Vanella, R., Boult, S., Küng, C. & Nash, M. A. Decoding the substrate specificity landscape of a promiscuous enzyme through multi-substrate mutational scanning. Nat. Commun. 17, (2026).

43. Yang, B. et al. Engineering the mechanical stability of a therapeutic complex between Affibody and Programmed Death-Ligand 1 by Anchor Point selection. ACS Nano 18, 31912–31922 (2024).

44. Sun, Y. et al. Engineering mechanostable Anticalin scaffolds to enhance particle adhesion and targeting of CTLA-4 under shear stress. Angew. Chem. Int. Ed Engl. 64, e202504483 (2025).

45. Walsh-Korb, Z., et al. Probing the scalability of ultra stable catch bond complexes. bioRxiv (2026) doi:10.64898/2026.01.08.695900.

46. Willaert, R. G. Adhesins of yeasts: Protein structure and interactions. J. Fungi (Basel) 4, E119 (2018).

47. Wrenbeck, E. E. et al. Plasmid-based one-pot saturation mutagenesis. Nat. Methods 13, 928–930 (2016).

48. Wong, E. et al. The Vibrio cholerae colonization factor GbpA possesses a modular structure that governs binding to different host surfaces. PLoS Pathog. 8, (2012).

49. Huang, X. et al. Vibrio cholerae biofilms use modular adhesins with glycan-targeting and nonspecific surface binding domains for colonization. Nat. Commun. 14, 2104–2104 (2023).

50. L, Kavanaugh Nicole, Q, Zhang Angela, J, Nobile Clarissa, D, Johnson Alexander & Katharina, R. Mucins Suppress Virulence Traits of Candida albicans. MBio 5, 10.1128/mbio.01911-14 (2014).

51. Valle Arevalo, A. & Nobile, C. J. Interactions of microorganisms with host mucins: a focus on Candida albicans. FEMS Microbiol. Rev. 44, 645–654 (2020).

52. Takagi, J. et al. Mucin O-glycans are natural inhibitors of Candida albicans pathogenicity. Nat. Chem. Biol. 18, 762–773 (2022).

53. Nett, J. E. et al. Host contributions to construction of three device-associated Candida albicans biofilms. Infect. Immun. 83, 4630–4638 (2015).

54. Gbejuade, H. O., Lovering, A. M. & Webb, J. C. The role of microbial biofilms in prosthetic joint infections. Acta Orthop. 86, 147–158 (2015).

55. Neoh, K. G., Li, M., Kang, E.-T., Chiong, E. & Tambyah, P. A. Surface modification strategies for combating catheter-related complications: recent advances and challenges. J. Mater. Chem. B Mater. Biol. Med. 5, 2045–2067 (2017).

56. Wilkinson, H. N. & Hardman, M. J. Wound healing: cellular mechanisms and pathological outcomes. Open Biol. 10, 200223 (2020).

