## Supplementary Information for "The *Candida auris* Adhesin Scf1 Mediates Broad Host-Protein Recognition through Shared Adhesive Regions"

This file contains:
Supplementary text: Materials and Methods
Supplementary Figures 1 to 5
Supplementary Tables 1 and 2
References

#### Supplementary text: Materials and Methods

*Native and recombinant human protein ligands used in this study:* Recombinant and native forms of putative human protein ligands used in this study were purchased from commercial vendors. Sources and catalog numbers are as follows: MUC1 (*Antibodies Online*, cat. no. ABIN1543605), VIT (*Sigma-Aldrich*, cat. no. 5051), FN (*Sigma-Aldrich*, cat. no. F0895), fibrinogen (*Sigma-Aldrich*, cat. no. F3879), human collagen type I (*Sigma-Aldrich*, cat. no. CC050), human collagen type IV (*Sigma-Aldrich*, cat. no. CC076), His-tagged MUC1 (*Acro Biosystems*, cat. no. MU1-H52H9), His-tagged VIT (*SinoBiological*, cat. no. 10424-H08H), His-tagged FN (*SinoBiological*, cat. no. 10314-H08H) and His-tagged albumin (*Acro Biosystems*, cat. no. HSA-H5220).

*Cloning*: The gene corresponding to the N-terminal domain of Scf1 from *Candida auris* strain B8441 (UniProt ID: A0A2H1A319; amino acids 28–255) was obtained from Twist Bioscience and cloned into the pCHA yeast expression vector using Gibson assembly.[1] The Scf1 gene had an N-terminal HA tag, allowing detection of expression using fluorescent antibodies. Plasmids were transformed into NEB 5-alpha chemically competent cells. The Scf1 gene and protein sequences are shown in the Supplementary Information (**Tables S1** and **S2**). Positive clones were identified by Sanger sequencing (Microsynth), and plasmids were extracted using the GeneJET Plasmid Miniprep Kit (Thermo Fisher Scientific).

*Yeast transformation and expression*: Plasmids were transformed into the EBY100 strain (Saccharomyces cerevisiae), which is auxotrophic for tryptophan due to a deletion of a tryptophan-biosynthesis gene. The EBY100 strain carries a genomic insertion encoding the Aga1 mating agglutinin protein, which is regulated by a galactose promoter. This enabled galactose-inducible expression of Scf1 fused to Aga2, which was then displayed on the yeast surface via a disulfide linkage between Aga1 (anchored to the cell wall) and Aga2 (encoded on the plasmid used for expression) (**Fig. 1B**). The resulting topology of the protein displayed on the yeast surface was as follows (N' to C'): *HA-Scf1-Myc-Aga2-yeast cell surface*, which mimics the natural configuration of Scf1 on the surface of *C. auris*. The pCHA plasmid carries a TRP1 selectable marker, allowing selection of transformants on agar plates with synthetic drop-out medium lacking tryptophan and containing 10% glucose. A single colony was transferred into liquid drop-out medium and grown overnight at 30 °C, 180 rpm. After 24 h, cells were transferred to a fresh liquid drop-out medium containing 1 M sodium phosphate buffer (pH 7) and 20% galactose for induction of protein expression to a final OD of ~0.5. Cells were grown for 24 h at 20 °C with shaking at 180 rpm. Cells were centrifuged at 13,000 g for 1 min and stored in phosphate-buffered saline with 0.1% bovine serum albumin (hereafter referred to as PBS-BSA). Protein expression levels were quantified by fluorescent antibody staining of the N-terminal HA tag. To detect expression, Two million yeast cells were centrifuged, the supernatant was discarded, and the cells were incubated with anti-HA primary antibody (1:500 dilution) for 30 min at room temperature. Cells were centrifuged at 13,000 RCF for 1 min and the supernatant was discarded. Cells were washed again with PBS-BSA to remove traces of unbound primary antibody. Cells were incubated with anti-mouse Alexa Fluor 594 secondary antibody (1:500 dilution) on ice for 20 min, followed by washing with PBS-BSA. Labeled cells were tested for expression using an Attune flow cytometer.

*Surface preparation and protein immobilization for spinning disk assay:* Circular cover glasses (25 mm diameter) were soaked in piranha etching solution (1:1 $H_2O_2$:sulfuric acid, 300 mL total volume) for 45 min, washed with distilled water and then incubated with epoxysilane (ABCR GmbH, Karlsruhe, Germany) on a rotary shaker for 1 h to functionalize the surface with reactive epoxy/glycidoxy groups, which allow covalent coupling of proteins and other biomolecules through amino-, thiol-, and hydroxyl-reactive chemistry. The cover glasses were rinsed three times with 100% ethanol, twice with Milli-Q water and then dried in an 80 °C oven for 1 h, cooled and stored under argon until further use. Various commercially available host-associated proteins diluted to 200 nM in 50 mM HEPES buffer, pH 8, were functionalized on the epoxysilane cover glasses using a sandwich method: a 90 µL drop of ligand was added to one slide and a second cover glass was placed onto it to allow the liquid to spread between the two glasses. Cover glasses were incubated for 1 h at RT, then washed three times with water. A solution of 90 µL PBS with 5% BSA was added to the cover glasses to block non-specific binding sites and incubated for 30 min at RT. To ensure that variation in ligand adhesion was not caused by inconsistent epoxysilane functionalization, all ligands were evaluated using the same batch of functionalized cover glasses. Furthermore, each non-binder experimental run (collagen I, collagen IV, and fibrinogen) included a known binder (e.g., MUC1) as a positive control to verify surface functionality.

*Spinning-disk adhesion assay:* Yeast cells expressing Scf1 were resuspended in 1 mL TBS buffer supplemented with 0.1% BSA and 1 mM calcium chloride, hereafter referred to as TBS buffer, and seeded on the surface of protein-functionalized cover glasses, followed by gentle shaking to achieve an even initial distribution over the surface. Yeast cells were incubated at room temperature for 1 h to adhere before spinning. The cell suspension was gently aspirated and replaced with TBS buffer. To calculate the number of cells before spinning, the cover glass was raster scanned and imaged frame by frame at 10× magnification (UPLFLN10X2) using bright-field settings on an Olympus IX81 microscope; approximately 500 individual images were automatically stitched together with CellSens software (version 1.16; Olympus, Tokyo, Japan) and saved in TIFF format. The buffer on the cover glasses was then removed gently by pipetting, and the cover glasses were mounted on the spinning disk device built in the Nash group, secured by a threaded metal holder and immersed in TBS buffer at room temperature. The spinning routine consisted of an acceleration ramp of 100 rpm/s, 5 min steady spinning at 3000 rpm, and a deceleration ramp of 100 rpm/s. During the adhesion assay, the cover glasses were maintained at a height of 25 mm from the bottom of the chamber to minimize turbulence and avoid disrupting the laminar boundary layer. The shear stress ($\tau$; dyn/cm$^2$) at any point on the surface of the cover glasses varies linearly with radial distance. After spinning, the cover glasses were raster scanned and imaged as described earlier to determine the number of cells that remained adherent after spinning. The percentage of cells retained on the cover glass was calculated relative to the number of cells on the cover glass before spinning. In parallel, the same process was carried out for Scf1(-) yeasts for each ligand.

Image analysis to generate cell-distribution heatmaps was performed using Python scripts. The yeast cells were segmented from the background by gray-value thresholding. The coordinates of the yeast cells were acquired as TSV files. The fraction of adherent yeast cells at different positions on the disk was calculated by normalizing the density of yeast cells (f) at each section of the disk to the density of yeast cells at the center of the disk, where shear stress is close to zero. The detachment profiles were plotted (f vs. $\tau$) and fitted with a sigmoid probabilistic model: $f = a/(1 + \exp[b(\tau - \tau 50)])$, where $\tau 50$ is the shear stress at which 50% of the cells remain adherent. This value was used as a measure of mean adhesion strength for comparison of yeast cell populations. To better view the distribution of cells, a density map was created with a Python script. The images were separated into square-shaped bins of the same size based on the coordinates. The number of cells within each bin was counted and displayed as a 2D heatmap.

*FACS-based binding assay*: Yeast cells expressing Scf1 were collected and stored in a PBS buffer (pH 7) supplemented with 0.1% BSA. A total of $10^6$ cells were incubated for 1 h with varying concentrations of His-tagged ligands (MUC1, VIT, FN and albumin) in TBS buffer (pH 7.2). Cells were centrifuged at 13,000 rpm for 1 min, and the supernatant was discarded. Then, 100 µL of mouse anti-His primary antibody (1:500) was added and samples were incubated for an additional 30 min. Samples were centrifuged at 13,000 rpm for 1 min, the supernatant was discarded, and cells were incubated with fluorescent Alexa Fluor 594 anti-mouse secondary antibody (1:500) for 30 min on ice. Cells were centrifuged, washed, and resuspended in TBS calcium buffer. The gated population was analyzed using an Attune flow cytometer, and median fluorescence values were recorded. To minimize variability in fluorescence signals arising from differences in Scf1 surface expression between expression rounds, all three ligands were tested in parallel for each independently induced Scf1-expressing yeast culture. To determine the apparent Kd values for each ligand, the median fluorescence values from replicate ligand-binding measurements were first averaged at each ligand concentration, and the standard error of the mean (SEM) was calculated across replicates. For each ligand, the raw fluorescence was fitted as a function of ligand concentration using Hill's binding model:

$$Y = \mathrm{Bottom} + \mathrm{Span} \times [L]^{nH} / (K_{D,app}{}^{nH} + [L]^{nH})$$

where $Y$ is the raw fluorescence signal, $[L]$ is the ligand concentration in µM, $Bottom$ is the fitted baseline signal, $Span$ is the fitted binding-associated signal increase, $K_{D,app}$ is the apparent dissociation constant, and $n_H$ is the Hill coefficient. Fitting was performed by nonlinear least-squares regression using a custom Python script.

***Deep mutational scanning of Scf1 ligand binding***

Deep mutational scanning experiments of the N-terminal Scf1 region were performed using protocols adapted from Vanella et al.[2,3] The gene encoding the N-terminal adhesive domain of *C. auris* Scf1 (residues 28–255 of the full-length protein) was introduced into the pUC19 plasmid, followed by whole-plasmid amplification using NNN mutagenic primers in accordance with the standard one-pot nicking mutagenesis protocol described previously.[4]

*PacBio barcoding and Scf1 library construction:* For DMS analysis, these residues were renumbered as construct positions 1–228. Following mutagenesis, unique DNA barcodes, hereafter referred to as unique molecular identifiers (UMIs), were linked by PCR to the mutagenized Scf1 inserts using the method described by Vanella et al.[1] The barcoded Scf1 inserts were then cloned into the pCHA yeast-display vector in the same configuration as the wild-type construct and propagated as a pooled plasmid library in *Escherichia coli*. Plasmid DNA extracted from the pooled bacterial library was subjected to Pacific Biosciences (PacBio) long-read sequencing before transformation into yeast. Because each PacBio read encompassed both the Scf1 coding sequence and its associated barcode, the sequencing data were used to generate a barcode–variant lookup table linking each UMI to the corresponding Scf1 sequence. The PacBio-characterized pCHA plasmid library was subsequently transformed into *Saccharomyces cerevisiae* EBY100 as described earlier. The resulting pooled library displayed the Scf1 variants as Aga2 fusion proteins at the yeast cell surface as follows (N′ to C′): HA–Mutant Scf1–Myc–Aga2–yeast cell surface.

*Sorting of expression and ligand-binding populations*: The surface expression of the mutant library was induced using galactose as described earlier. The induced library ($40 \times 10^6$ yeast cells) was stained for the N-terminal HA epitope using mouse anti-HA primary antibody and Alexa Fluor 594-conjugated goat anti-mouse secondary antibody under the same conditions described earlier. For the ligand-binding selections, independent aliquots of $40 \times 10^6$ yeast cells were incubated for 1 h at room temperature with 200 nM MUC1-His, 20 nM VIT-His, or 80 nM FN-His in TBS containing 0.1% BSA. These concentrations were chosen to be near their apparent Kd values (Fig. 3). Bound ligand was detected by incubation with mouse anti-His primary antibody at a 1:500 dilution for 30 min, followed by Alexa Fluor 594-conjugated goat anti-mouse secondary antibody at a 1:500 dilution for 30 min on ice. The fluorescence-shifted populations from the expression and ligand-binding samples were sorted using a FACS Melody cell sorter into four bins, numbered 1–4, spanning low to high fluorescence (**Fig. S3**). Sorted cells were pooled according to their bins and grown for 24–48 h at 30 °C in -TRP glucose medium until the OD reached 6–9, equivalent to 60–90 million cells, and stored at −80 °C as glycerol stocks. All expression and binding experiments were performed in two independent biological replicates.

*DNA preparation and sequencing:* Plasmid DNA from glycerol stocks was isolated using Zymo Yeast Miniprep Kit. The mutagenized Scf1 coding region was amplified using primers containing sample-specific Illumina index sequences. Amplicons were purified, pooled at equimolar concentrations, and sequenced using Illumina sequencing. Sequencing reads were quality-filtered, assigned to their corresponding sorting bin and biological replicate, and aligned to the wild-type Scf1 sequence. Reads containing insertions, deletions, mutations outside the targeted region, or multiple amino-acid substitutions were excluded. The number of reads corresponding to each Scf1 variant was tabulated separately for each bin.

*Calculation of Scf1 expression and ligand-binding fitness scores*: Illumina sequencing reads assigned to each Scf1 variant were converted to estimated cell counts within each fluorescence-activated cell sorting bin. For variant $v$, the cell count $c_v$was calculated from its fraction of total sequencing reads:

$$\frac{r_v}{r_{tot}} = \frac{c_v}{c_{tot}},$$

where $r_v$ and $r_{tot}$ are the variant-specific and total read counts, respectively, and $c_v$ and $c_{tot}$ are the corresponding variant-specific and total sorted-cell counts. A fluorescence-weighted mean, $\beta_v$, was then calculated across all sorting bins:

$$\beta_v = \frac{\sum_{i=1}^{n_{bins}} \omega_i c_{vi}}{\sum_{i=1}^{n_{bins}} c_{vi}},$$

where $c_{vi}$ is the estimated number of cells containing variant $v$ in bin $i$, and $\omega_i$ is the median fluorescence intensity of that bin. Expression and ligand-binding fitness scores were calculated by normalizing the weighted mean of each variant to that of wild-type Scf1:

$$F = log_2\left(\frac{\beta_v}{\beta_{WT}}\right)$$

Because substitutions that alter surface expression can also influence binding, the MUC1-, VIT-, and FN-binding fitness scores were normalized to the corresponding expression fitness score. The fitness scores were first transformed into non-logarithmic values and the expression-normalized binding score for variant $v$ and ligand $L$ was calculated as:

$$2^{F_{v,L}^{norm}} = \frac{2^{F_{v,L}^{bind}}}{2^{F_v^{expression}}}$$

where $L$ denotes MUC1, VIT, or FN

Expression-normalized scores close to one indicate wild-type-like ligand binding after correction for surface-display level. Values less than one indicate a specific loss of ligand binding that cannot be explained solely by reduced Scf1 expression, whereas values greater than one indicate enhanced binding relative to the amount of displayed protein. The final expression-normalized substitution scores for each variant at each position were displayed as position-by-amino-acid heatmaps (**Fig. 4A, main text**). The binding and sorting experiments were carried out in two biological replicates, which showed good correlation (**Fig. S4).**

*Calculation of Position- and Patch-Level Fitness Reduction for Scf1 Ligand Binding:* The analysis was performed independently for MUC1, VIT, and FN using a custom Python script. Synonymous substitutions and missing fitness measurements were excluded. For each ligand, the median fitness reduction at position $p$ was calculated across all measured non-synonymous substitutions:

$$R_p = 1 - median(F_{p,a})$$

where $F_{p,a}$ is the normalized fitness of substitution $a$ at position $p$, and a fitness score of 1 represents WT-like activity.

The fraction of substitutions below WT at each position was calculated as:

$$B_p = \frac{number\ of\ measured\ substitutions\ with\ F_{p,a} < 1}{total\ number\ of\ measured\ substitutions\ at\ position\ p}$$

For each predefined patch $P$, the script calculated the arithmetic mean of the available position-level values:

$$R_P = \frac{1}{n_P}\sum_{p\in P} R_p\ ;\ B_P = \frac{1}{n_P}\sum_{p\in P} B_p$$

where $n_P$ is the number of evaluable positions within patch $P$.

The non-patch background was calculated by averaging the corresponding position-level values across all evaluable positions outside the five predefined patches:

$$R_{\text{background}} = \frac{1}{n_Q}\sum_{p\in Q} R_p\ ;B_{\text{background}} = \frac{1}{n_Q}\sum_{p\in Q} B_p$$

where $Q$ denotes the set of evaluable positions outside the predefined patches, and $n_Q$ is the number of positions in $Q$.

*Single-clone validation:* Single-mutant Scf1 clones used for DMS-validation experiments (**Fig. 4C, main text**) were generated by site-directed mutagenesis at selected positions in the Scf1 gene. Cloning, expression, and ligand-binding (DMS-validation) experiments were performed as described above. All single-mutant clones showed robust surface expression comparable to wild-type Scf1, indicating that the mutations did not abolish yeast-surface display (**Fig. S5**). Fitness scores for single mutants were calculated in the same manner as for the DMS library and normalized for differences in expression level.

*Structure Prediction and Visualization*: The N-terminal Scf1 protein structure was predicted using AlphaFold2. The resulting model was visualized, colored, and modified in PyMOL to generate the final structural representation.

*Statistical calculations:* Statistical comparisons for **Fig. 2** (main text) were performed using two-sided Welch's unpaired t-tests. Selected comparisons were made between ligand groups under the Scf1 condition (i.e., MUC1 vs. VIT, MUC1 vs. FN, etc.), and within each ligand between Scf1 and Scf1(-) yeasts. Data are presented as mean ± SEM. For **Fig. 4** (main text), Pearson's correlation coefficient and p-values (two-tailed) were calculated by comparing DMS-derived and single-clone fitness scores. Significance was defined as $p < 0.0001$ (****), $p < 0.001$ (***), $p < 0.01$ (**), $p < 0.05$ (*) and $p \geq 0.05$ (n.s).

### SUPPLEMENTARY FIGURES

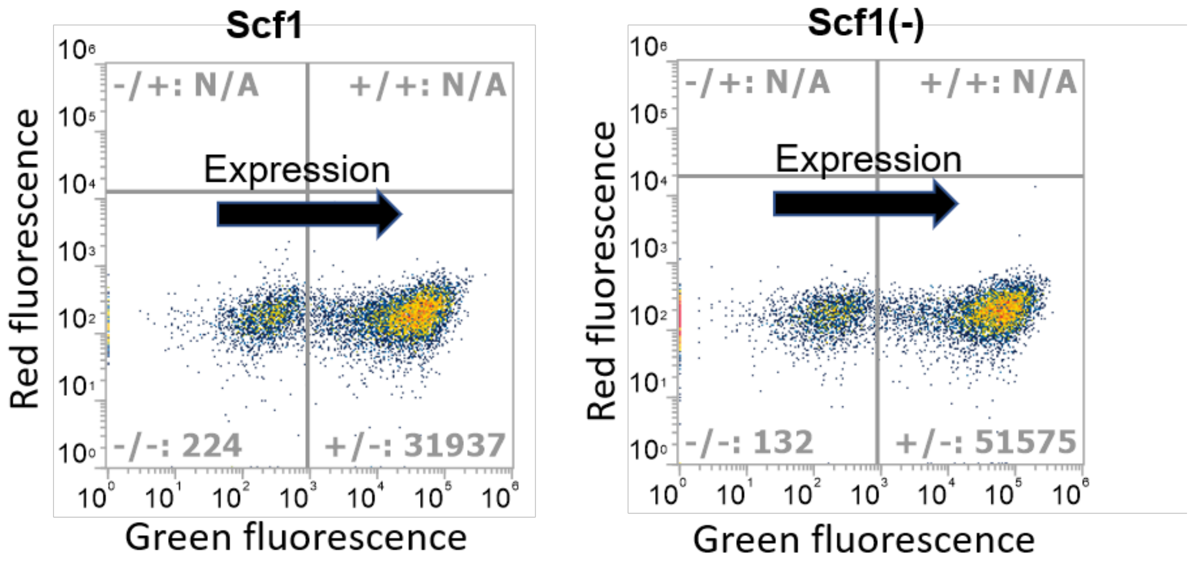


**Figure S1: Flow cytometry-based detection of Scf1 display on yeast**. Cell density plots demonstrate galactose-induced surface expression of Scf1 and Scf1(-) at 24 h, detected by a fluorescent antibody against the N-terminal HA tag. Each dot represents a single yeast cell. The y-axis shows red fluorescence, while the x-axis shows green fluorescence. Warmer colors indicate higher event density. The arrow denotes increasing expression. Grey numbers indicate event counts within the displayed gates/quadrants; “N/A” indicates quadrants for which no events were reported.

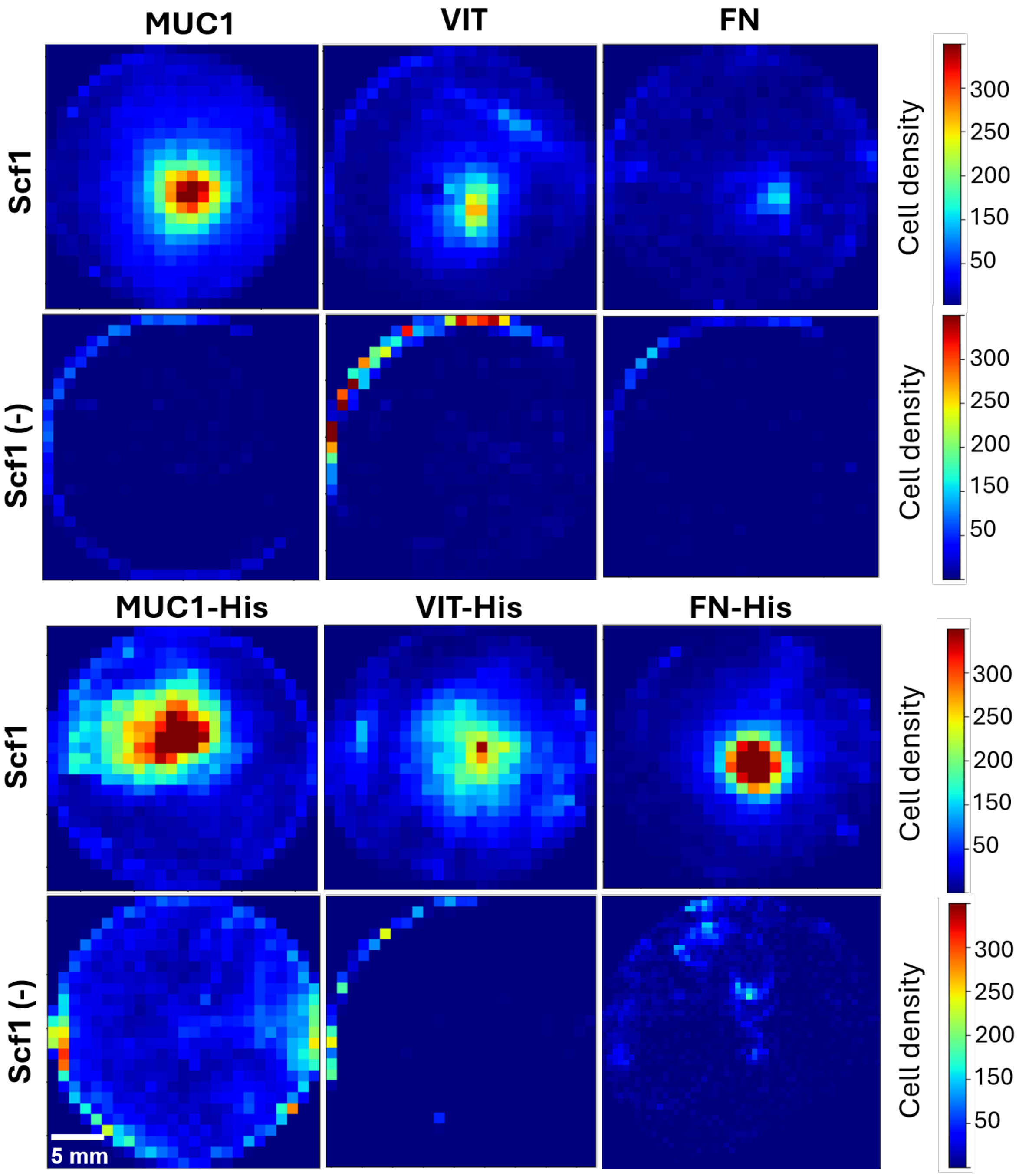


**Figure S2: Comparison of adhesion of Scf1 and Scf1(-) to ECM ligands under shear stress**. Heatmaps show the spatial density of yeast cells displaying Scf1 or Scf1(-) that remained attached to 25-mm cover glasses coated with different human protein ligands after spinning (3000 rpm, 5 min). For visualization, each post-spin image was partitioned into 25 × 25–pixel bins. Cells in each bin were counted and mapped to a colormap. The color scale shows the number of detected cells per spatial bin with warmer colors indicating higher local cell density (0–300 cells/bin; 5-mm scale bar in the lower left). The rotation axis is typically at the center of each panel, and shear stress increases with radius. Panels show superimposed (averaged) heatmaps from two to three independent experiments for the indicated ligands.

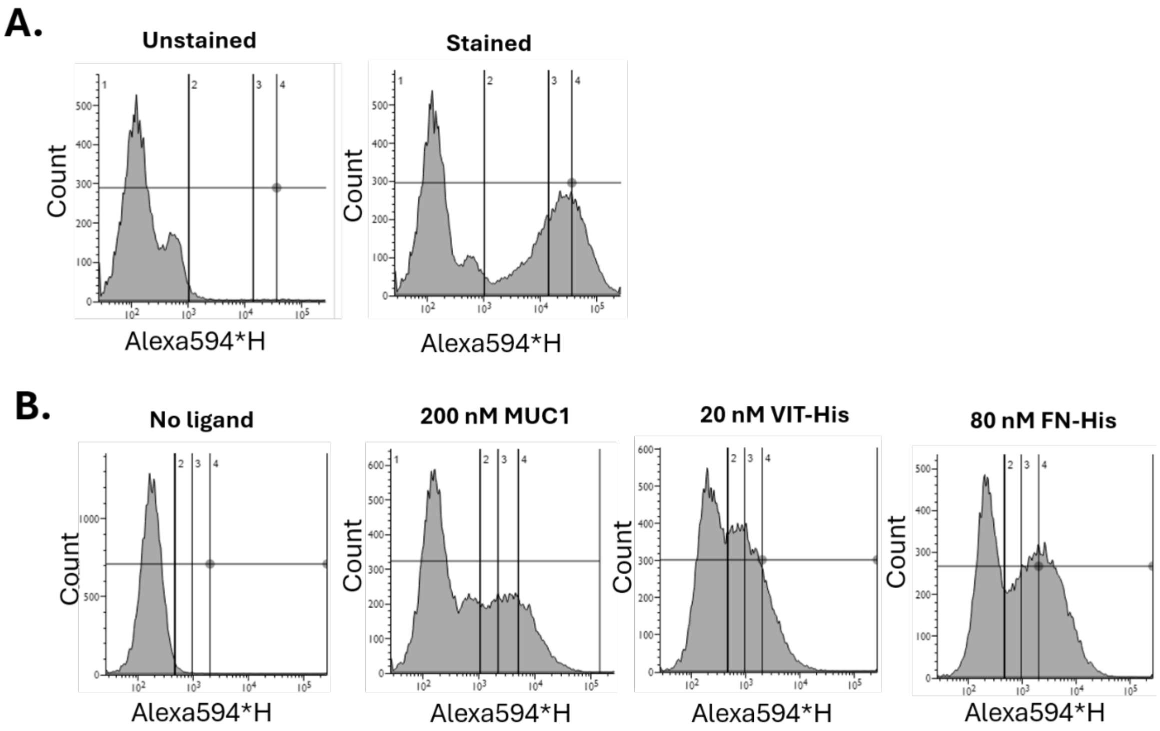


**Figure S3: Flow-cytometric sorting of the Scf1 deep-mutational-scanning library according to surface expression and ligand binding.** Shown here are representative fluorescence histograms of singlet-gated yeast cells displaying the mutagenized Scf1 library. **Top panel:** surface-expression staining. Surface expression was detected using a mouse anti-HA primary antibody followed by an Alexa Fluor 594-conjugated anti-mouse secondary antibody. **Bottom panel:** ligand-binding selections. From left to right, histograms show the no-ligand control and libraries incubated with 200 nM MUC1-His, 20 nM VIT-His, or 80 nM FN-His. Bound ligand was detected using a mouse anti-His primary antibody followed by an Alexa Fluor 594-conjugated anti-mouse secondary antibody. Histograms show event count as a function of Alexa Fluor 594 fluorescence intensity (Alexa594-H). Vertical boundaries divide each population into four sorting bins, numbered 1–4 from low to high fluorescence, representing progressively increasing Scf1 surface expression or ligand-binding signal. Cells collected from each bin were expanded separately and used for downstream sequencing and fitness-score calculation. Expression and ligand-binding selections were performed in two independent biological replicates.

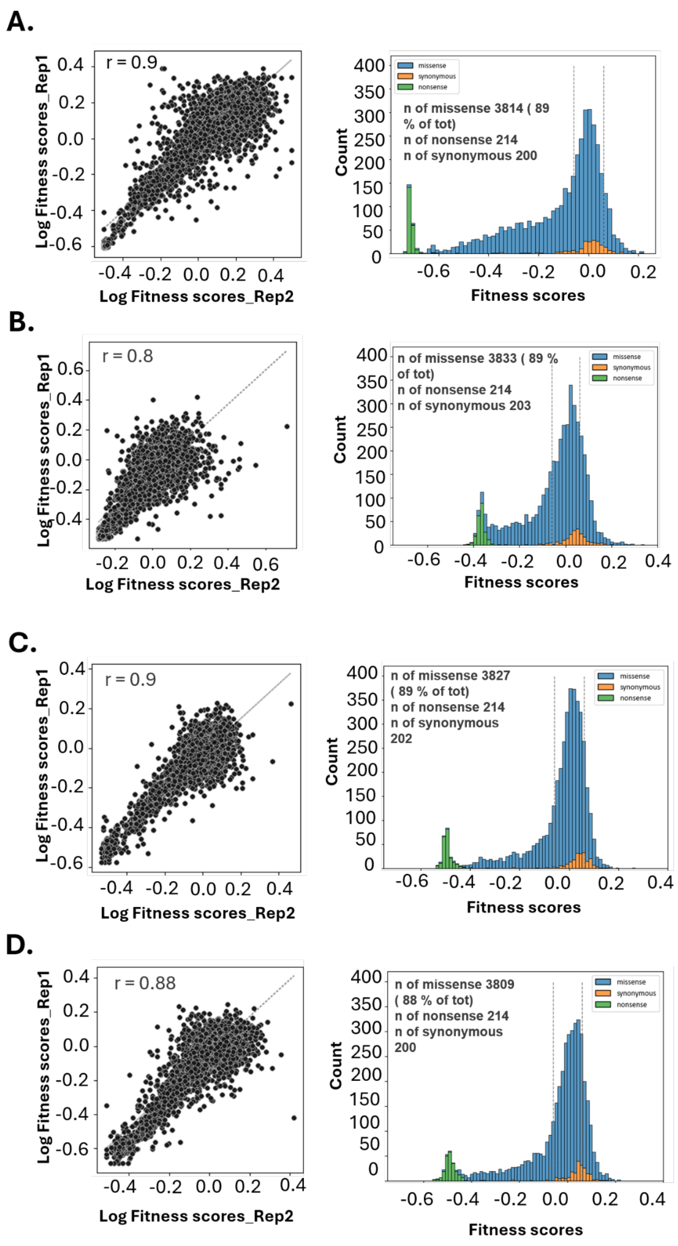


**Figure S4: Reproducibility and fitness-score distributions of Scf1 deep mutational scanning datasets.** Scatter plots show the correlation between variant fitness scores obtained from two independent biological replicates for **A.** Scf1 library surface expression, **B.** MUC1-binding, **C.** VIT-binding, and **D.** FN-binding. Each point represents an individual Scf1 sequence variant detected in both biological replicates. Dashed diagonal lines indicate the line of identity, and the corresponding Pearson correlation coefficients (*r*) are shown in each panel. Histograms show the distributions of fitness scores obtained for each DMS dataset, with variants classified as missense, synonymous, or nonsense substitutions. Dashed vertical lines indicate the fitness-score range defined by synonymous variants and therefore provide an experimental reference for WT-like fitness, following the approach described by Vanella et al.[1] Missense substitutions (blue) show a broader distribution of fitness effects than synonymous substitutions (orange), whereas nonsense substitutions (green) are predominantly shifted toward reduced fitness. For the ligand-binding datasets, fitness scores represent MUC1-, VIT-, or FN-binding fitness measured by FACS-based sorting and sequencing. Numbers of detected missense, synonymous, and nonsense variants are indicated within the respective panels.

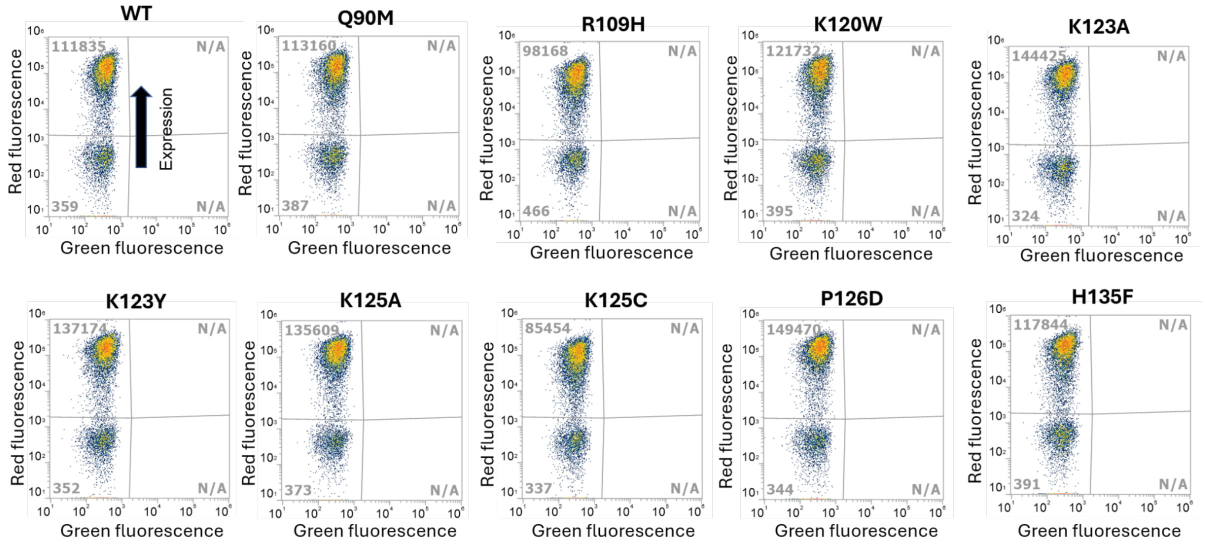


**Figure S5: Surface-expression analysis of individually reconstructed Scf1 variants**. Representative flow-cytometry density plots showing surface expression of wild-type Scf1 and selected single-substitution variants reconstructed from the DMS dataset. Yeast cells displaying each Scf1 construct were stained for the N-terminal HA epitope using a mouse anti-HA primary antibody followed by an Alexa Fluor 594-conjugated anti-mouse secondary antibody, as described in the Supplementary Methods. Each dot represents an individual yeast cell, with warmer colors indicating higher event density. The vertical red fluorescence shift indicates increased Scf1 surface expression. Grey numbers indicate event counts within the displayed gates/quadrants; "N/A" denotes quadrants for which no events were reported. The analyzed variants show a clear HA-positive population comparable to wild-type Scf1, indicating that the reconstructed substitutions do not abolish yeast-surface display of the Scf1 construct.

### SUPPLEMENTARY TABLES

**Table S1: Gene sequences used in this study**

| Gene | Gene sequence |
|---|---|
| Scf1 | **TACCCATACGACGTTCCAGACTACGCT**ATTGGTACTACCACATTACCTGCTGATGATG<br>ATTACTGTACCCCATCTAAACATTATTGGGTTAAAGGTGGTCACCATAGAAGATCAAA<br>TGATGGTCCAGAATTTATTGATGCAGGTGTGGCGGTTTCCGACGTTACTAAAGTTGGT<br>GATAATGTTTACGAGTTGTCTTTGAACTTTAAAACAGCTGAAGATGAAGAATTGGCTA<br>AAGCTATTTCTAACAATCAAGTTAGATCTCTGACTATTGAAGGTACTGGTGTTGAAGA<br>CGTCCAATTGATTGGAAGAGATAACAAAATGGAGGGTATTCCATGGACAAAATGGACA<br>GCTAGAGTTAGAGTTAGGTCTGAAAGAATGTTGGAAAAAAGACATAAACATAAGCCAG<br>CTGGTGTAGTCTGCTGTTTACCACATAAGTTGGAGGTAAGATTGGCTTTGGAACCAGA<br>AGGTGAAGCAGCTAGAGCTTACAAGAGCGTTTTTTCTAGAGACTCATACGCCTATACT<br>GTGGAAAGAGGTGTTGGTGATGAAATTACCGAAGTTGGTGACGTTGATGGTTTTTTGA<br>ATAATGTGCTAAAGAGAGACGTTGATTCTCCTTTGTCAAAAAGATCCGGTGTCTTCAA<br>TAGAGTTAGAAATTTGCAACCAAGATTCCATAAGCAATGTTGGGTTTGTGAATGTGAA<br>ACCACCACTACTACAGGCTCCGGATCCACGCGTGCA**GAACAAAAGCTTATCTCCGAAG**<br>**AAGACTTG**CAGGAACTGACAACTATATGCGAGCAAATCCCCTCACCAACTTTAGAATC<br>GACGCCGTACTCTTTGTCAACGACTACTATTTTGGCCAACGGGAAGGCAATGCAAGGA<br>GTTTTTGAATATTACAAATCAGTAACGTTTGTCAGTAATTGCGGTTCTCACCCCTCAA<br>CAACTAGCAAAGGCAGCCCCATAAACACACAGTATGTTTTTTAA |
| Scf1(-) | **TACCCATACGACGTTCCAGACTACGCT**TGCACGCGTGGATCCAGAACCGCCCACATTA<br>ACACCACCATCAAAAAACTGGGTTTTCTGTTCAACCAGATCGTTATTGGCAAAGCCAC<br>CAACTTCAACAAATTTGATAACGCTCAGACCATTCGGTGCACCACTTTTAACTTTTGC<br>CACAATGGTTGCAAAAACACCATCTTCGGTAATTGCATAGGCACCTGTACCGCTATCT<br>TCGGCAAACAGAAAAACAATCATTTTGCGATCCGGATAAACTGCGGTATCAAAGCTTT<br>TGGTCGGATTCGGATCAACGATCAGTTCACCAGGTTCAATTTCAATGATTTCCAGCAC<br>GTTCGGATCATAGCTATAAACGAAATCGCAATTGGCAATACCTTTGCTCGGAATACCG<br>CTAAAACGAACCGGAATACGCACGGTATCACCCGGTTTTGCATTAACGGTATCCACTT<br>TAATACGAACGGCATCCAAATCAGTGCTAGC**GAACAAAAGCTTATCTCCGAAGAAGAC**<br>**TTGCAGGAACTGACAACTATATGCGAGCAAATCCCCTCACCAACTTTAGAATCGACGC**<br>**CGTACTCTTTGTCAACGACTACTATTTTGGCCAACGGGAAGGCAATGCAAGGAGTTTT**<br>**TGAATATTACAAATCAGTAACGTTTGTCAGTAATTGCGGTTCTCACCCCTCAACAACT**<br>**AGCAAAGGCAGCCCCATAAACACACAGTATGTTTTTTAA** |

The HA tag is shown in bold black; the NScf1 gene sequence is shown in red, followed by a short linker sequence. The c-Myc tag is shown in bold blue, and the sequence encoding the Aga2p mating factor is shown in bold green. The NScf1(–) construct contains the same vector backbone, yeast display scaffold, Aga2p fusion, and flanking epitope tags, HA and c-Myc, but encodes an unrelated cohesin domain of CipA protein from the bacterium *Clostridium thermocellum* [5] instead of Scf1 (shown in magenta). It therefore served as a negative control in the yeast-surface-display experiments. This design ensures that any differences in adhesion can be attributed solely to the displayed NScf1 rather than vector or display artifacts.

**Table S2: Protein sequences**

| Protein | Protein sequence (N- to C-terminal topology) |
|---|---|
| Scf1 | **YPYDVPDYAIGTTTLPADDDYCTPSKHYWVKGGHHRRSNDGPEFIDAGVAVSDVTKVGDNVYELSLNFKTAEDEELAKAISNNQVRSLTIEGTGVEDVQLIGRDNKMEGIPWTKWTARVRVRSERMLEKRHKHKPAGVVCCLPHKLEVRLALEPEGEAARAYKSVFSRDSYAYTVERGVGDEITEVGDVDGFLNNVLKRDVDSPLSKRSGVFNRVRNLQPRFHKQCWVCECETTTTT**GSGSTRA**EQKLISEEDLQELTTICEQIPSPTLESTPYSLSTTTILANGKAMQGVFEYYKSVTFVSNCGSHPSTTSKGSPINTQYVF*** |
| Scf1(-) | **YPYDVPDYA**ASTDLDAVRIKVDTVNAKPGDTVRIPVRFSGIPSKGIANCDFVYSYDPNVLEIIEIEPGELIVDPNPTKSFDTAVYPDRKMIVFLFAEDSGTGAYAITEDGVFATIVAKVKSGAPNGLSVIKFVEVGGFANNDLVEQKTQFFDGGVNVGGSGSTRA**EQKLISEEDLQELTTICEQIPSPTLESTPYSLSTTTILANGKAMQGVFEYYKSVTFVSNCGSHPSTTSKGSPINTQYVF*** |

Protein sequences are shown in their N- to C-terminal topology with color-coded functional elements. In the Scf1 construct, the HA tag is shown in bold black at the N-terminus, followed by the Scf1 sequence in red, a short linker sequence, the c-Myc tag in bold blue, and the C-terminal Aga2p fusion in bold green. The Scf1(–) control retains the same N- to C-terminal arrangement and color-coded display architecture but contains an unrelated sequence corresponding to the seventh cohesin domain of the CipA protein [4] (shown in magenta) in place of Scf1 and served as a negative control (Fig. 2, main text). The Scf1(-) yeasts showed significantly lower adhesion to MUC1, VIT, and FN than NScf1 yeasts, indicating that increased cell retention is mediated by the Scf1 sequence rather than nonspecific retention by the display scaffold.

#### References


1. Gibson, D. G. *et al.* Enzymatic assembly of DNA molecules up to several hundred kilobases. *Nat. Methods* **6**, 343–345 (2009).
2. Vanella, R. *et al.* Understanding activity-stability tradeoffs in biocatalysts by enzyme proximity sequencing. *Nat. Commun.* **15**, 1807 (2024).
3. Vanella, R., Boult, S., Küng, C. & Nash, M. A. Decoding the substrate specificity landscape of a promiscuous enzyme through multi-substrate mutational scanning. *Nat. Commun.* **17**, (2026).
4. Wrenbeck, E. E. *et al.* Plasmid-based one-pot saturation mutagenesis. *Nat. Methods* **13**, 928–930 (2016).
5. Stahl, S. W. *et al.* Single-molecule dissection of the high-affinity cohesin-dockerin complex. *Proc. Natl. Acad. Sci. U. S. A.* **109**, 20431–20436 (2012).
